# Metabolic similarity and the predictability of microbial community assembly

**DOI:** 10.1101/2023.10.25.564019

**Authors:** Jean C.C. Vila, Joshua Goldford, Sylvie Estrela, Djordje Bajic, Alicia Sanchez-Gorostiaga, Alejandro Damian-Serrano, Nanxi Lu, Robert Marsland, Maria Rebolleda-Gomez, Pankaj Mehta, Alvaro Sanchez

## Abstract

When microbial communities form, their composition is shaped by selective pressures imposed by the environment. Can we predict which communities will assemble under different environmental conditions? Here, we hypothesize that quantitative similarities in metabolic traits across metabolically similar environments lead to predictable similarities in community composition. To that end, we measured the growth rate and by-product profile of a library of proteobacterial strains in a large number of single nutrient environments. We found that growth rates and secretion profiles were positively correlated across environments when the supplied substrate was metabolically similar. By analyzing hundreds of in-vitro communities experimentally assembled in an array of different synthetic environments, we then show that metabolically similar substrates select for taxonomically similar communities. These findings lead us to propose and then validate a comparative approach for quantitatively predicting the effects of novel substrates on the composition of complex microbial consortia.

## Introduction

Quantitatively predicting shifts in the genetic and phenotypic composition of populations in response to environmental change is a major challenge in ecology and evolution (Lässig, Mustonen, and Walczak 2017; Wortel et al. 2023; Cadotte et al. 2015). In microbial systems, there is widespread interest in predicting the effects of different metabolites on community composition and function. For example, one would like to predict how dietary compounds impact the composition of the gut microbiome (Human Microbiome Project Consortium 2012; Faith et al. 2011; David et al. 2014; Wu et al. 2011), how different fertilizers alter the composition of soil communities (Allison, Hanson, and Treseder 2007; Sha et al. 2023), and how different metabolites determine the composition of industrial microbial consortia (Lindemann et al. 2016; Roell et al. 2019). Quantitative tools for predicting how microbial community composition responds to shifts in the metabolic environment will be critical if we wish to engineer and control their composition and direct these communities towards desirable states (Sanchez et al. 2023; Chacón and Harcombe 2019). However, due to the complexity and variability of species-species and species-environment interactions, it is unclear if the impact of different metabolites on the composition of complex microbial communities is even predictable (Chacón and Harcombe 2019; van den Berg et al. 2022).

Over the past few years, our lab and others have used multi-replicated enrichment communities as a model system to study the repeatability of community assembly across environments (Goldford et al. 2018; Dal Bello et al. 2021; Aranda-Díaz et al. 2023). These in-vitro systems allow one to quantify reproducible similarities and differences in community composition when one systematically manipulates individual components of the metabolic environment one at a time (Estrela, Sánchez, and Rebolleda-Gómez 2021; Estrela et al. 2021). A major discovery from this body of work is that microbial communities assembled in identical metabolic environments converge to reproducible compositions at coarse phylogenetic and functional levels, despite variability in species composition (Goldford et al. 2018; Shan et al. 2023). In parallel, similar patterns of compositional variability and reproducibility have been observed in many naturally assembled microbiomes, ranging from the mouse gut (Turnbaugh et al. 2009) to the open ocean (Louca, Parfrey, and Doebeli 2016).

In nutrient limited environments, the reproducibility of community assembly arises from selection on metabolic traits, e.g., growth rates or by-product secretion (Estrela, Vila, et al. 2022).

Because these traits are quantitatively similar in phylogenetically related taxa, related species can fulfill equivalent functional roles leading to communities with similar coarse-grain phylogenetic compositions despite differences at the species level (Martiny et al. 2015; Aguirre de Cárcer 2019; Estrela, Vila, et al. 2022). Additionally, recent studies involving high-throughput measurements of microbial growth phenotypes and pairwise inter-specific interactions in different environments have shown that different environments can sometimes be ’coarse-grained’ and grouped together based on the chemical class of substrate present (Tong et al. 2020; Kehe et al. 2021; Gralka, Pollak, and Cordero 2022). These two observations suggest that similar initial metabolic environments may lead to quantitative similarities in metabolic traits, which we reasoned would lead to selection for quantitatively similar communities. In this paper we therefore set out to experimentally test the hypothesis that the composition of microbial communities assembled in one metabolic environment may be predictable by observing the composition in a quantitatively similar metabolic environment (Fig 1A).

**Figure 1:**
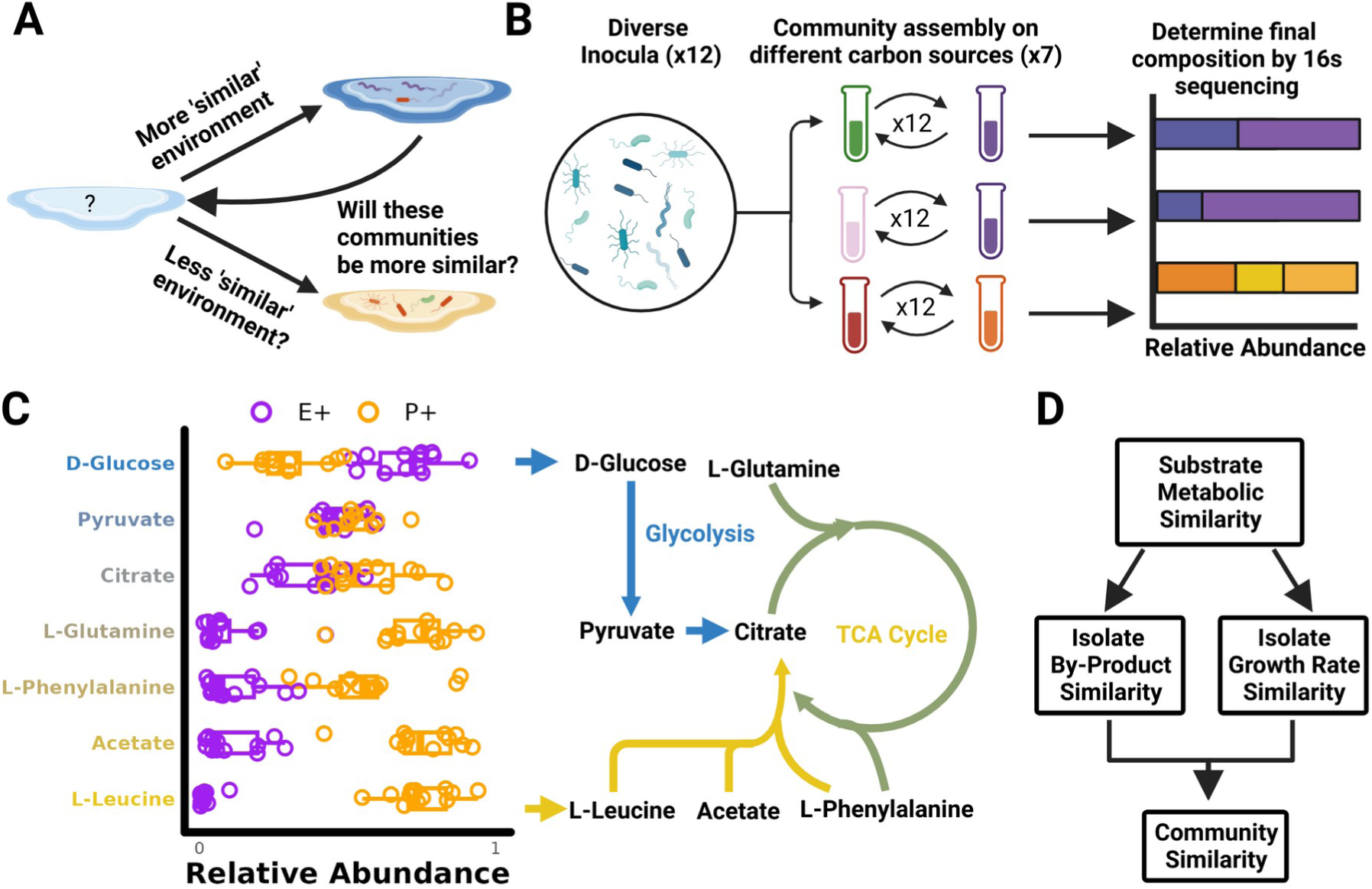
Is microbial community assembly predictable across similar environments? A) Our goal is to determine whether one can predict the effect of novel environments on the composition of diverse microbial consortia. We hypothesize that community composition in new environments might be predictable using the community composition observed in similar metabolic environments (B) To test this hypothesis, we turned to an in vitro community system in which diverse inocula are assembled in minimal media containing different carbon sources. We assembled enrichment communities from 12 inocula by passaging for 12 transfers on one of 7 different carbon sources. (C) In this panel we show the relative abundance of the two dominant proteobacterial clades after 12 transfers (Methods). The purple E+ clade includes *Enterobacteriaceae*, *Aeromonadaceae*, *Erwiniaceae*, and *Yersianiaceae*, while the orange P+ clade includes *Pseudomonadaceae* and *Moraxellaceae*. Different carbon sources resulted in different community compositions at this coarse-grain phylogenetic level with a shift in composition that appears to qualitatively track a primary axis in central metabolism. Arrows indicate the entry point of D-Glucose (Blue) and L-Leucine (Yellow) into central metabolism (D) Given this observation, we hypothesize that microbial isolates grown on substrates using similar metabolic pathways would exhibit similarities in growth rate and secrete similar metabolic by-products. If our hypothesis is true, then we expect metabolically similar substrates to select for predictably similar microbial communities.

### Community composition depends on the metabolism of the supplied substrate

In our previous work we assembled communities from different inoculum in minimal media containing different carbon sources and showed that they converge to reproducible compositions at coarse phylogenetic levels (family level or higher), (Goldford et al 2018). We have additionally provided a mechanistic explanation for the composition of communities assembled on d-glucose as the sole carbon source (Estrela, Vila, et al. 2022). On d-glucose, communities contained quantitatively similar abundances of two gammaproteobacterial clades (Fig. S1A). The most abundant fermenter clade (henceforth E+), including the Enterobacteriaceae, Aeromonadaceae, Erwiniaceae, and Yersianiaceae families, is selected for the ability to grow rapidly on d-glucose while secreting quantitatively similar levels of the metabolic by-product acetate (Fig. S1B-C). The second most abundant obligate respirator clade (henceforth P+), including the Pseudomonadaceae and Moraxellaceae families, is selected for the ability to grow rapidly on secreted organic acids (Fig. S1D).

Given the reproducibility of community assembly on d-glucose media, we first wanted to explore how shifting carbon sources would alter community composition. We therefore repeated our community assembly experiments, this time assembling enrichment communities from 12 different inocula under serial passaging on minimal media containing one of 7 different carbon sources (Fig 1B). In addition to d-glucose these carbon sources included other intermediates of glycolysis and the tricarboxylic acid (TCA) cycle (pyruvate, acetate and citrate) as well as amino acids (l-leucine, l-glutamine and l-phenylalanine) (see Methods). As in our previous work communities were grown at 30C without shaking and then diluted 125 fold with fresh media every 48 hours. At the end of 12 growth cycles (∼84 generations, a time we have recently shown is enough to reach a state of stable equilibrium (Chang et al. 2023)), communities on all 7 carbon sources are typically dominated by the same E+ and P+ gammaproteobacterial clades (Fig 1C) we had previously observed. However, these clades reached very different abundances depending on the carbon source. Specifically, we observed a shift from E+ to P+ dominated communities that appears to track a primary axis of central metabolism, with carbon sources higher up in glycolysis selecting for higher levels of E+, and carbon sources entering at lower glycolysis and in the tricarboxylic acid (TCA) cycle selecting for higher levels of P+ (Fig 1C).

The striking observation that microbial community composition mirrors central metabolic pathways suggests that the structure of microbial metabolic networks might play a pivotal role in shaping how different substrates select for different communities (Dal Bello et al. 2021; Takano et al. 2023). As previously established on d-glucose, coarse-grain community composition is largely determined by the taxonomic distribution of growth rates and by the amount of secreted metabolic by-products (Fig S1) (Estrela, Vila, et al. 2022). We therefore hypothesized that, metabolically similar carbon sources processed by highly overlapping pathways (e.g. D-Glucose and D-Fructose), should exhibit similar growth rates, and also result in similar by-product profiles. In turn, we hypothesize that similarities in these key traits will result in similar community compositions when communities are assembled on similar substrates (Fig 1D). In the following sections we experimentally test these hypotheses.

### The taxonomic distribution of growth rates is similar on metabolically similar substrates

To determine whether similar environments lead to similar growth rates we turned to a library of proteobacterial strains we had previously isolated either from glucose enrichment communities or from environmental sources (Methods). From an initial library of 112 isolates we selected 62 strains that had unique full length 16s gene sequences available (Fig S2, Methods). This library covered the most abundant families we had found in our communities and included Enterobacteriaceae (n=35), Erwiniaceae(n=2),Yersiniaceae (N=2), Pseudomonadaceae (n=18), Aeromonadacae(n=1), Moraxellaceae(n=2) Comamonadaceae(n=1), and Alcaligenaceae (n=1). We measured the growth curves of all 62 strains over 48 hr in M9 minimal media supplemented with 1 of 19 different carbon sources (Fig2A, Methods). The 19 carbon sources were selected to span a range of chemical classes and included sugars, sugar alcohols, organic acids and amino acids. (Fig. S3). From each growth curve we measured the average growth rate of each strain on each carbon source (*R_iα_*) and then mapped this trait onto a 16s maximum likelihood phylogenetic tree. (Fig. 2B, Methods).

**Figure 2:**
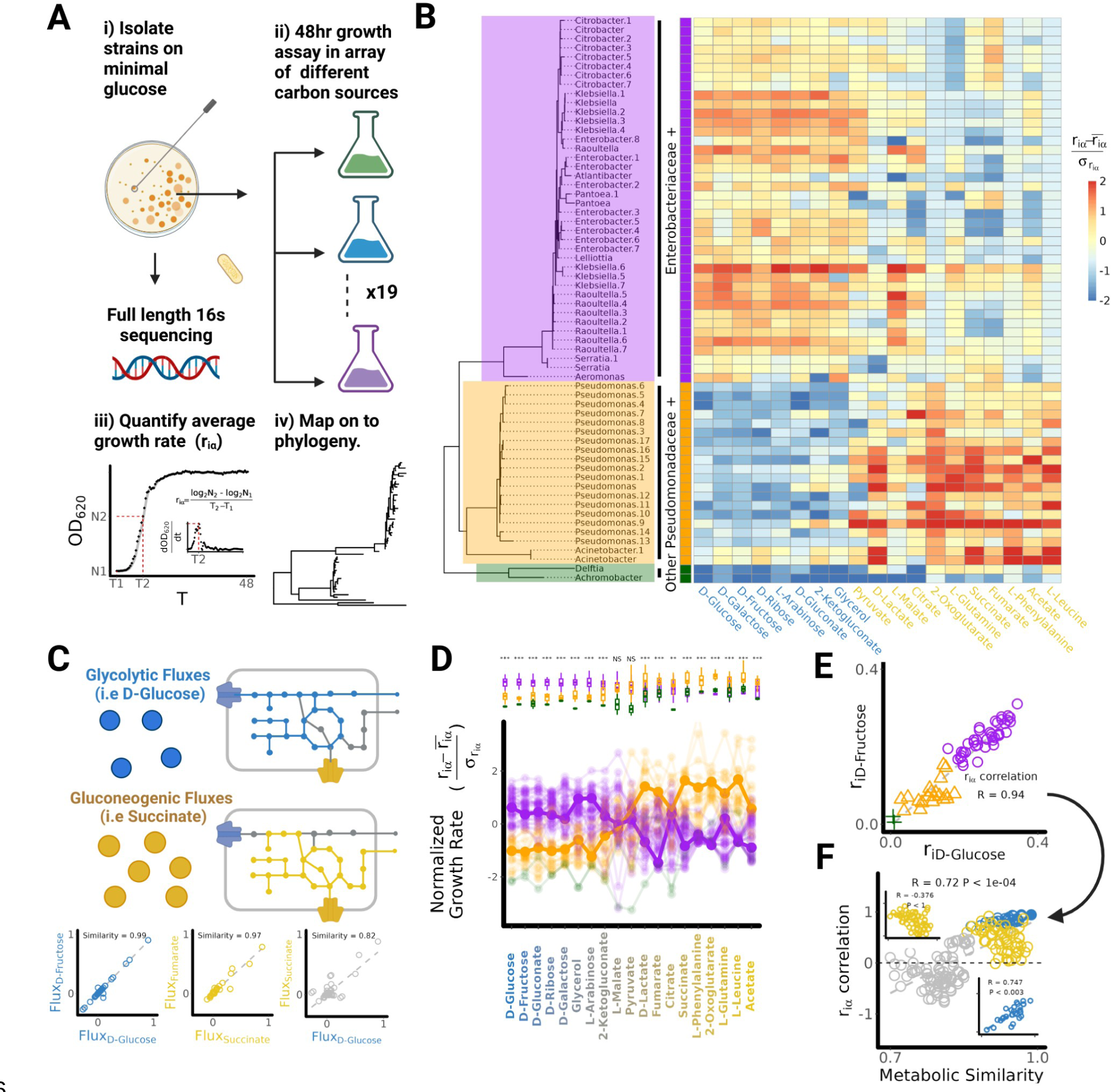
A quantitative measure of substrate similarity predicts growth rate similarities (A) Diagram of experimental procedure. 62 proteobacteria isolates with unique 16s sequences where isolated (Fig. S2) and grown in minimal media containing one of 19 different carbon sources. The average growth rate of each isolate on each carbon source was quantified and normalized to allow for comparison across environments (Methods). (B) Phylogenetic heat map shows the taxonomic distribution of growth rates. Carbon sources are ordered heuristically by the order of entry into central metabolism (Fig. S3). Carbon sources that enter metabolism via glycolysis are coloured in Blue (Glycolytic) and Carbon sources entering metabolism via the TCA cycle are coloured in yellow (Gluconeogenic). (C) To quantify the similarity of substrate pairs, we performed flux-balance analysis on individual carbon sources and calculated the correlation in predicted intracellular metabolic fluxes (methods). Sub-panels illustrate this calculation for D-glucose and D-Fructose which are both glycolytic, Fumarate and Succinate which are both gluconeogenic and D-Glucose and Succinate. (D) When we rank order carbon sources by quantitative similarity to D-Glucose, we find that all Enterobacteriaceae+ strains (purple) have higher growth rates on carbon sources that are more quantitatively similar to D- Glucose whereas all Pseudomonadaceae+ strains (orange) have higher growth rates on carbon sources that are more dissimilar to D-Glucose. (E) Quantitatively similar carbon sources such as D-Glucose and D-Fructose display a strong positive correlation in growth rate (see Fig S7 for correlation between all pairs of carbon sources). (F) More positive growth rate correlations are observed between more metabolically similar carbon sources (Mantel Test P<1e-04). Stronger positive correlations are observed for pairs of more similar glycolytic resources (blue inset), but not for more similar gluconeogenic carbon sources (yellow inset). The weakest correlations are observed when comparing growth on a glycolytic carbon source to growth on a gluconeogenic carbon source (gray circles, (pearson’s r = -0.172±0.34, mean±sd)).

Consistent with our initial hypothesis we found that on d-glucose and other sugars (i.e d-fructose, d-galactose, d-ribose and l-arabinose), the taxonomic distributions of growth rate were highly similar (Fig.2B). Specifically, on all five sugars, members of the E+ clade grow faster than members of the P+ clade (p ≤ 0.0001, two sample T-test) (Fig 2B, Fig. S4A), which was also true for other carbon sources including non-sugars, e.g. d-gluconate, 2-ketogluconate, and glycerol (p ≤ 0.0001, two sample T-test). Within the E+ clade members, the *Klebsiella* and *Raoultella* genera had higher growth rates compared to other genera (p ≤ 0.0001, two sample T-test, Fig. S4B). Recent studies have classified carbon sources based on whether they enter metabolism via glycolysis (glycolytic) or the TCA cycle (gluconeogenic) (Gralka et al 2022, Borer et al 2022). When we classify our 19 carbon sources along this binary, we find that all eight carbon sources on which E+ grew significantly faster than P+ were glycolytic, whereas all nine carbon sources on which P+ grew significantly faster than E+ were gluconeogenic (Fig. S5). Notably, no significant differences were observed between the E+ and P+ clades on pyruvate and l-malate, which are metabolic intermediates between glycolysis and the TCA cycle (Sauer and Eikmanns 2005). The above experiment thus confirms that substrates using the same metabolic pathway tend to have a similar taxonomic distribution of growth rates. This observation led us to hypothesize that carbon sources metabolized using quantitatively more similar pathways would exhibit more correlated growth rates.

### A quantitative measure of substrate similarity predicts growth rate similarities

In order to test this hypothesis we need to define a quantitative measure of substrate similarity. For this purpose, we turned to flux balance analysis (FBA), a widely used metabolic modeling approach that predicts metabolic fluxes in different environments (Orth, Thiele, and Palsson 2010; Chacón and Harcombe 2019). We constructed a modified version of the *E. coli* genome-scale metabolic model iJO1366 by supplementing it with a minimal set of additional reactions ensuring growth on every carbon source considered in this study (Methods, Table S1) (Orth et al. 2011). We then introduced a metric of metabolic similarity for each pair of carbon sources defined as the Pearson’s correlation coefficient of the intracellular metabolic fluxes predicted by FBA for the optimal metabolism of each (Fig. 2C, Methods). Whilst the model used to quantify substrate similarity is based on *E. coli*, metabolic networks are broadly conserved in heterotrophic microbes. As such, substrate similarities calculated using metabolic models from different organisms tend to be highly correlated to the substrate similarities calculated with the modified iJO1366 model (Fig. S6) Using this metric of metabolic similarity, we could rank-order all carbon sources by their metabolic similarity to d-glucose (Fig. 2D). As shown in Fig. 2D, all 40 E+ strains exhibit an average decline in their normalized growth rate as the carbon source became increasingly dissimilar to d-glucose (Spearman’s ⍴ = -0.65 ±0.13 (Mean ±SD)). Conversely all 20 P+ strains exhibited an average increase in normalized growth rate as the carbon source became more dissimilar to d-glucose (Spearman’s ⍴ = 0.78 ±0.07 (Mean ±SD)). To quantify similarities in growth rates across environments, we calculated the Pearson correlation between the growth rate of all isolates in one carbon source vs another. A representative example is shown in Figure 2E for d-glucose and d-fructose. For this pair of highly similar carbon sources, we observed a strong positive correlation in growth rates for our isolate library (R=0.94, two-sample correlation test p<0.001). We then extended the same procedure to determine the correlation between growth rates in every other pair of carbon sources (Fig. S7), plotting it against our computationally predicted measure of their metabolic similarity (Fig. 2F). We found that growth rates are indeed more positively correlated on more metabolically similar carbon sources (R=0.72, Mantel test p<0.001). Interestingly, stronger growth rate correlations are observed on pairs of more similar glycolytic carbon sources (R=0.747, Mantel test p<0.003) but not on pairs of more similar gluconeogenic carbon sources (R=-0.76, Mantel test p=1.0) (Fig. 2F, insets). The correlation between metabolic similarity and growth rate correlation is robust to alternative measures of growth rate (Methods, Fig. S7). These results illustrate that our measure of metabolic similarity is able to both predict the previously described binary resource classification as well as capture some of the subtle differences in growth rate distributions across different environments.

### Metabolically similar substrates lead to the secretion of similar by-products

We have demonstrated that the distribution of growth rates is more similar on more metabolically similar carbon sources (Fig 2). However, even in an environment containing a single limiting resource, community composition is not solely determined by growth on supplied resources; secretions and consumption of metabolic by-products play a critical role (Goldford et al., 2018). Because closely related taxa tend to use similar metabolic pathways, they also tend to secrete similar metabolic by-products (Borenstein et al., 2008; Plat et al., 2015; Han et al., 2022). For example in our isolate library we have conducted targeted measurements of the amount of 5 by-products released after 16 hours of growth on d-glucose (see Methods). These by-products were chosen because they were known to be released by two different pathways for glucose metabolism: acetate, lactate and succinate are secreted by mixed acid fermentation (Vivijs et al. 2015) whereas gluconate and 2-ketogluconate are released by periplasmic glucose oxidation (Molina et al. 2019). Because of these different metabolisms, members of the E+ clade secrete acetate, d-lactate and succinate whereas members of the P+ clade secrete gluconate and 2-ketogluconate (Fig. S9). Just as different strains secrete similar by-products because they use similar metabolic pathways, we reasoned that different substrates using similar metabolic pathways should lead to the secretion of similar by-products.

In order to test whether secretions on other glycolytic carbon sources would be similar to those observed on d-Glucose, we conducted an experiment using four different strains – an Enterobacter and Pseudomonas strain from one of our glucose enrichment communities (C2R4), as well as two reference laboratory strains belonging to the E+ and P+ clades: *E. coli MG1655* and *P. putida KT244*. These four strains were inoculated into minimal media containing either d-glucose or one of four other carbon sources (n = 2 biological replicates). Two of these carbon sources were glycolytic (d-fructose, and glycerol), while two were gluconeogenic (l-malate and pyruvate). We selected these four carbon sources so that all four strains grew to detectable densities (OD600 >0.04) whereas this was not true for most other carbon sources.

After 16, 28, and 48 hours of growth, we collected the spent media of each strain on each carbon source and characterized the secretion profile using targeted liquid chromatography– mass spectrometry (LC-MS) (Methods). We also repeated this experiment for the spent media on 5 other carbon sources for a subset of timepoints and strains (Methods). Isotope-labeled internal standards allowed us to quantify the absolute concentration of up to 143 metabolites of which 20 were detected across all samples.

In Fig. 3B, we plot the concentration of the three most abundant secreted by-products (acetate, lactate and succinate) as a function of time when the *Enterobacter* strain is grown on each of the five carbon sources. The full dataset, which includes the other strains and carbon sources and the accumulation dynamics for 17 additional by-products is shown in Fig. S10. For all carbon sources and strains we observe numerous by-products after 16 hours of growth. By 48 hrs the concentration of secondary metabolites decreases indicating a switch from consumption of supplied resources to consumption of secreted by-products (Wang et al., 2021). To determine whether similar carbon sources led to similar by-products, we calculated the similarity in by- product profiles across carbon sources using the Pearson’s correlation in metabolite abundance at each time point. Consistent with our hypothesis, we find that after 16 hours of growth, by- products on the glycolytic carbon sources (d-fructose and glycerol) were highly similar to those on d-glucose for all 4 strains and in both replicates (Pearson’s R = 0.93±0.11 (mean±sd, N=16)). In contrast we observed more variable by-product correlations to D-Glucose on gluconeogenic carbon sources (l-malate and pyruvate) (Pearson’s R = 0.3 ±0.41, mean±sd, N=16). This difference held when we repeated the analysis for the 28 and 48 hour time points, though we observed less variability in secretion profile across carbon sources potentially due to the confounding effect of tertiary by-products produced from the consumption of secondary by- products (Fig. S11).

**Fig 3:**
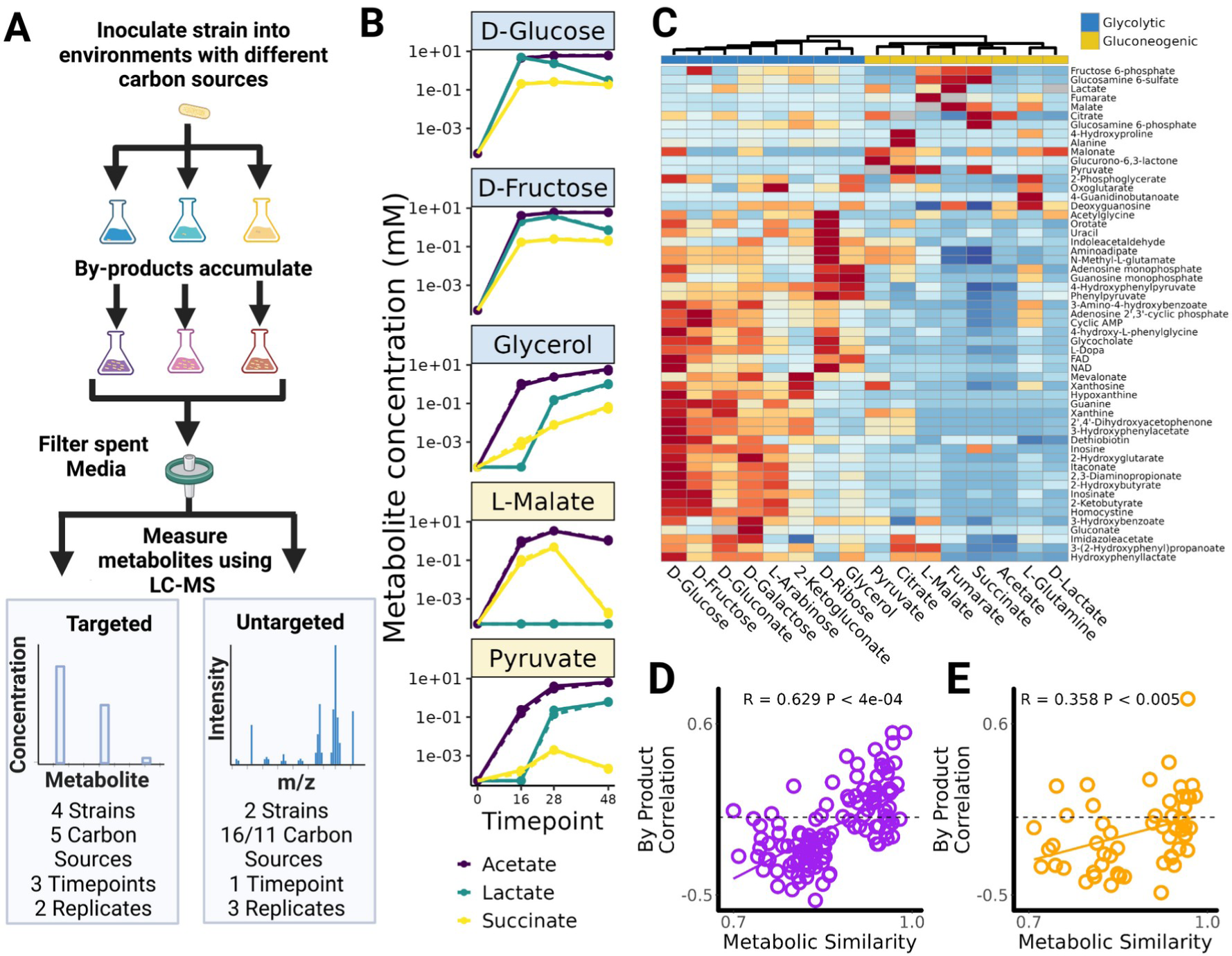
Metabolic secretions are similar in metabolically similar environments. (A) Diagram of experimental procedure. Strains were inoculated and grown on minimal media supplemented with different carbon sources. Samples were filtered to obtain the spent media and the metabolite profile was quantified using either targeted or untargeted metabolomics (LC-MS). Targeted LC-MS was used to quantify absolute concentration of a known set of metabolites at multiple time-points. Untargeted LC-MS was used to compare the metabolite profile across many carbon sources at a single time-point. (B) Concentrations of the 3 most abundant metabolites secreted by an *Enterobacter* strain on 5 different carbon sources (see Fig S10 for other metabolites and strain). Similar secretions are observed when strains are grown on similar carbon sources such as D-Glucose and D-Fructose (Fig. S11). (C) Untargeted LC-MS on the spent media of a *Klebsiella* strain after 24hrs of growth on 16 different carbon sources. Heatmap shows normalized peak height for each metabolite averaged across 3 replicates (see Fig. S13A for individual replicates). Red cells correspond to carbon sources on which each metabolite is more abundant whereas blue cells correspond to carbon sources on which each metabolite is less abundant. Carbon sources are clustered based on the correlation in by- product profile. (D) *Klebsiella* has a more correlated by-product profile when grown on more similar carbon sources (Mantel Test, P<2e-04). (E) A *Pseudomonas* strain grown on 11 carbon sources displays a qualitatively similar relationship (Mantel Test, P<0.005) (Fig. S12,S13B)

Building on this first experiment, we next asked whether there existed a positive relationship between metabolic similarity and the correlation in metabolic by-product profiles. To obtain sufficient statistical power for this analysis, we conducted a second metabolomics experiment involving a new pair of strains, this time growing the strains on a wider array of carbon sources and with the goal of identifying a larger number of secretions. We selected a representative *Klebsiella* (E+), and *Pseudomonas* (P+) strain and grew them on all 19 carbon sources previously studied as well as in a no-carbon source control (n= 3 replicates). We collected the spent media after 24 hours of growth and performed untargeted metabolomics (LC-MS) on all carbon sources for which significant growth had been detected in all three replicates (OD600 >0.04). This gave us 16 carbon sources on which the *Klebsiella* strain grew sufficiently and 11 carbon sources on which the *Pseudomonas* strain grew sufficiently. We detected and annotated 69 peaks across all samples, and focused on the 55 metabolites that were absent from the no-carbon-source control (Methods).

By comparing the peak heights of these 55 metabolites across all samples, we can quantify relative changes in by-product secretion when consuming different substrates. In Figure 3C we show the normalized peak height for all 55 secretions for the *Klebsiella* strain averaged across 3 biological replicates (Methods). We clustered carbon sources based on the correlation in by- product profile (Pearson’s correlation in normalized peak height) and found that, for the *Klebsiella* strain, glycolytic carbon sources cluster together as do the gluconeogenic carbon sources (Fig. 3C). For the *Pseudomonas* strain the 3 glycolytic carbon sources cluster together as do 7 out of the 8 gluconeogenic carbon sources (Fig S12). Qualitatively similar results are observed at the individual replicate level (Fig S13). Using our metabolic similarity metric (Fig 2C) we can test the hypothesis that quantitatively similar carbon sources had more positively correlated secretion profiles. To that end, we calculate the by-product correlation (Pearson’s R) for every carbon source pair and find that, for the *Klebsiella* strain, more metabolically similar carbon sources tended to lead to more correlated by-product profiles (R=0.629, Mantel test p<2e-04) (Fig 3D). A qualitatively similar though weaker relationship is observed for the *Pseudomonas* strain (R=0.358, Mantel test p<0.005) (Fig. 3E). These correlations hold when we analyze each of the three individual replicates independently rather than averaging across replicates (Fig. S14). These results confirm that, for individual strains, more metabolically similar substrates will typically lead to more similar metabolic by-products.

### Metabolically similar substrates select for taxonomically similar communities

In sum, we have provided evidence that a quantitative measure of substrate similarity based on flux balance predicts similarities in growth rates (Fig. 2) and similarities in secreted by-products (Fig. 3). In our previous work we had found that community composition on D-Glucose was well predicted by the growth on the supplied substrate as well as on the secreted by-products (Goldford et al. 2018; Estrela, Vila, et al. 2022). We therefore hypothesized that our quantitative measure of substrate similarity will predict similarities in community composition.

To test this hypothesis, we needed to analyze a large number of communities assembled in a large number of single-nutrient environments. Following a similar community assembly protocol to the experiment described in Figure 1C (Methods), we assembled serially passaged, enrichment communities in up to 42 different carbon sources from three different inocula (Fig 4A). In Figure 4B, we show the relative abundance of the E+ and P+ clades after 70 generations (10 serial passages) in our library of carbon sources (see Fig S15 for exact sequence variant (ESV) level composition). For all three inocula, the relative abundance of E+ decreases as carbon sources become increasingly dissimilar to d-glucose (Spearman’s ⍴ = -0.83 ±0.04 (Mean ±SD)) while the relative abundance of P+ increases (Spearman’s ⍴ = 0.64±0.09 (Mean ±SD)). This shift from E+ to P+ dominated communities matches the shift from E+ to P+ preferred carbon sources (Fig 2D). Specifically, E+ reaches a higher relative abundance on glycolytic carbon sources where the average growth rate of E+ strains is higher (Pearson’s R = 0.94 ±0.03 (Mean ±SD)), while P+ reaches higher relative abundance on gluconeogenic carbon sources where the average growth rate of P+ strains is higher (Pearson’s R =0.77 ±0.16 (Mean ±SD)) (Fig. S16). This result confirms that the previously identified similarities in growth rate and by-product secretions observed from individual strains extend to the community level (Fig. 2D).

**Fig 4:**
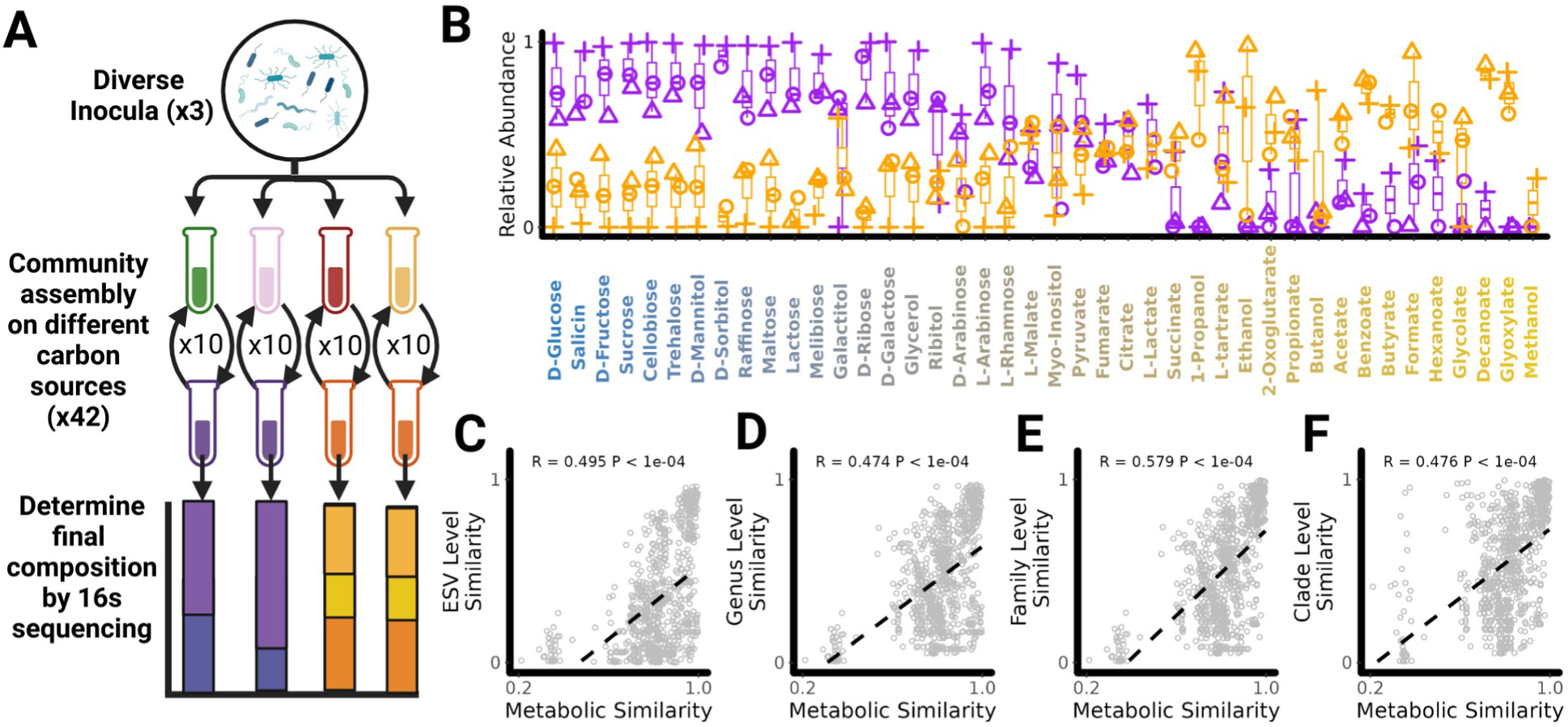
Metabolically similar carbon sources select for taxonomically similar communities. A) Diagram of experimental procedure. Diverse microbiomes were collected from 3 different environmental sources and used as inoculum for enrichment communities assembled in one of 42 different carbon sources. Communities were passaged every 48hrs for 10 transfers after which community composition was determined using 16s rRNA sequencing (Fig S15). (B) Relative abundance of the Enterobacteriaceae+ (Purple) and Pseudomonadaceae+ (Orange) clade on each carbon source after 10 transfers. Carbon sources are rank ordered along the X axis by the metabolic similarity to D-Glucose. Different shapes in this plot corresponds to communities assembled from different inocula. Metabolically similar carbon sources select for taxonomically similar communities at the (C) ESV level, (D), Genus level, (E) Family level, and (F) Clade level (i.e P+/E+). In Panels C-F we show this relationship for a single inoculum (see Fig. S17 for all three inocula). The correlation between metabolic similarity and community similarity was calculated for each inoculum separately and was statistically significant in every instance (Mantel Test P< 1e-4).

In order to quantify the relationship between metabolic similarity and community similarity, we calculated community similarities (Renkonen similarity) at the ESV, Genus, Family and “Clade” (E+/P+) level for every pair of communities assembled from each of the three inocula (Methods). Figures 4C-F show the relationship between metabolic similarity and community similarity for a single inoculum (see Fig S17 for all three). Consistent with our hypothesis, we found that in every inoculum and at every taxonomic scale considered, communities assembled on more metabolically similar substrates were more taxonomically similar (ESV-Level R=0.48± 0.02, Genus-Level R = 0.46 ± 0.01,Family-level R=0.56 ±0.02, Clade-Level R = 0.52±0.06, (mean±sd, N=3)). Consistent with previous findings from our group (Goldford et al. 2018; Estrela, Vila, et al. 2022), we found that communities assembled from different inocula have weaker correlations between metabolic similarity and community similarity at the ESV and Genus level compared to communities assembled from the same inoculum (Fig S18A-B). In contrast, much smaller differences are observed at the Family or Clade level (Fig S18C-D).

Despite this general pattern, we did observe systematic shifts in community composition across inocula even at higher levels of taxonomy (Figure 4B). For example, communities assembled from one of the three inocula were systematically enriched for E+ compared to the other two (median relative abundance: 0.63 vs 0.50 or 0.51 ; Mann-Whitney U test, p = 0.0123 or 0.007, one-tailed). In order to quantify how much of the variation in community composition was due to difference in inocula compared to differences in carbon source we turned to an additional data- set where we had assembled enrichment communities in one of 24 different carbon sources from eight different inocula (Fig S19A, Methods). The large amount of inocula in this data-set allowed us to decompose the total variation in community composition at different taxonomic levels (beta-diversity) into the variation due to the inocula and the variation due to the carbon source (Methods,(Legendre 2008)). We find that as we decrease the level of taxonomic resolution the proportion of variation explained by the inocula decreases from 15.1% at the ESV level to 5.6% at the Clade level (Fig S19B). In contrast the proportion of variation explained by the carbon source increases from 15.5% at the ESV level to 77% at the Clade level (Fig S19C) . This suggests that while metabolic similarity does correlate with community similarity the amount of predictable variation is limited at lower levels of taxonomic organization where differences in the initial species pool and other unexplained sources of variation play a larger role (Fig S19D) .

### Community assembly in new environments is predictable using a comparative approach

In the previous section we showed that metabolically similar substrates select for taxonomically similar communities (Fig 4). Specifically the strength of this relationship depended both on the taxonomic scale considered and the specific inocula examined with systematic shifts in community composition observed for different inocula (Fig 4B,S19). Given these results we reasoned that, in new environments, the fine-grain taxonomic structure of microbial communities would only be predictable when communities were assembled from the same species pool, whereas the coarse-grained taxonomic structure might be predictable even when communities are assembled from different species pools. If community structure is predictable how might one actually predict it? In protein homology modeling, the structure of a new ’target’ protein can be predicted by using the structure of a closely related ’template’ protein as determined by sequence similarity. By analogy, we propose that the simplest approach to predict community composition in a new ’target’ environment is to use the composition observed in the closest ‘template’ environment as determined by metabolic similarity (Fig 5A).

**Fig 5:**
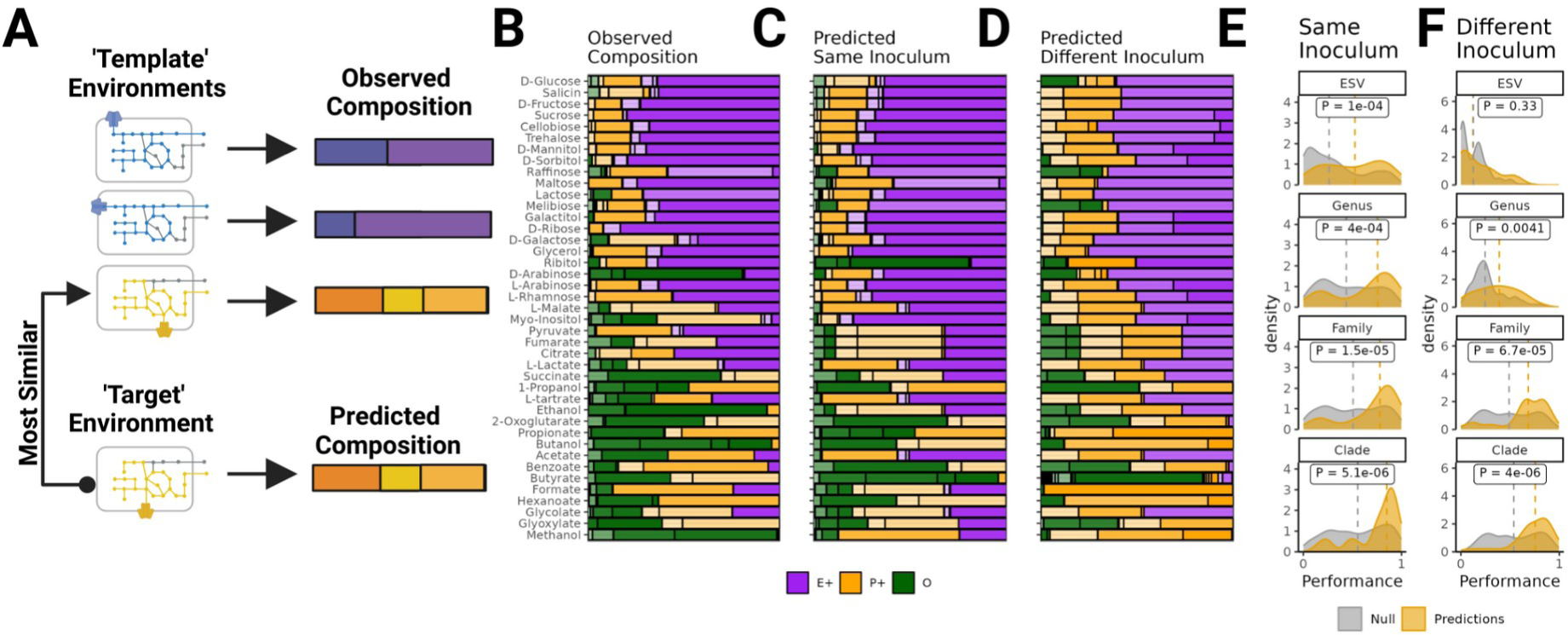
Community assembly in new environments is predictable using a comparative approach. (A) We propose a simple comparative approach for predicting community composition in new environments. The composition in a new ‘target’ environment can be predicted using community composition observed in the most metabolically similar ‘template’ environment. We evaluated the performance of this approach by conducting “Leave-One-Carbon-Source-Out” cross-validation using the communities assembled on different carbon sources (Fig 4). (B) Observed composition for communities assembled on different carbon sources (Inoculum 1). Different ESV are shown in different shades of Purple (E+ Strains), Orange (P+ strains) and Green (Other). (C) Composition predicted using the community assembled in the most metabolically similar environment from the same inoculum (Inoculum 1). (D) Composition predicted using the community assembled in the most metabolically similar environment from a different inoculum (Inoculum 2). The performance of these predictions is quantified by calculating the community similarity (Renkonen similarity) between predicted and observed composition on each carbon source at different taxonomic levels. (E) Distribution of performance for predictions made using the same inoculum (yellow) (F) Distribution of performance for predictions made using different inocula (yellow). We compared the performance of our comparative approach to the performance of a null model where predictions are made using a random environment (Grey). In Panels E-F, the dotted lines show the median performance for the prediction and the null. In each panel we test whether median performance using our comparative approach was higher than the performance of the null model using one tailed Mann-Whitney U-test

To evaluate the effectiveness of this comparative approach, we performed “leave-one carbon-source-out” analysis using the stable communities we had previously assembled from one of the three inocula (corresponding to the data shown in Fig. 4B) by passaging it on a library of different carbon sources (N=41) (Fig. 5B). For each carbon source, we predicted the community composition using the composition of the community assembled in the most metabolically similar environment, where metabolic similarity was determined based on modeled flux balance as described in the Methods. These predictions can be made using communities assembled from the same inoculum (Fig. 5C), as well as communities assembled from one of the other inocula (Fig. 5D). In order to quantify the prediction performance, we calculated the community similarity (Renkonen similarity) between the predicted and observed compositions on each carbon source at the ESV, Genus, Family and Clade level. A similarity of 1 indicates a perfect match between predicted and observed composition (identical relative abundances), whereas a value of 0 indicates no overlap between the predicted and observed composition (no shared taxa). In line with our expectations, predictions made using the same inoculum (Inoculum 1) on average performed significantly better than predictions made using a different inoculum (Inoculum 2) at the ESV level (median: 0.56 vs 0.11; Mann-Whitney U test, p < 8e-08, one-tailed) and Genus level (median: 0.76 vs 0.39; Mann-Whitney U test, test, p <1.3e-05,one-tailed). No significant differences were observed at the Family level (median: 0.79 vs 0.68; Mann-Whitney U test,p = 0.1, one-tailed) and Clade level (median: 0.85 vs 0.76; Mann-Whitney U test, p = 0.16, one-tailed).

In order to test whether predictions made using this comparative approach performed better than chance, we ran a permutation test where we compared the observed distribution of performances to a null distribution of performances where predictions are made using a random environment as the ‘template’ (Methods). For predictions made using the same inoculum (Inoculum 1), the comparative approach outperforms this null at all taxonomic resolutions considered (Fig. 5E, Mann-Whitney U test, p < 0.00039, one-tailed). In contrast, predictions made using a different inoculum (Inoculum 2) did not outperform the null model at the ESV level (Mann-Whitney U test,p = 0.39, one-tailed), performed marginally better at the Genus level (Mann-Whitney U test, p = 0.012, one-tailed), and significantly better at the Family or Clade level (p =4.7e-05 and p = 1.8e-05, Mann-Whitney U test, one-tailed).

To assess how general these observed patterns are, we repeated the above analysis for all possible combinations of predicted and observed inocula using data from all three assembly experiments we have analyzed in this paper, presented in Figures 1, 4 and S19 respectively (Fig. S20A-D,Methods). In total, this dataset includes 343 different communities assembled from 23 different inocula on 48 different carbon sources. When using the same inoculum the comparative approach performs consistently better than the null at all taxonomic levels considered (Fig. S20E). When using a different inoculum the comparative approach performed quantitatively similarly to the null at the ESV level, marginally better at the Genus Level and substantively better at the Family and Clade level (Fig. S20F). These results illustrate that a significant fraction of the variation in community compositions is predictable across metabolically similar environments; at a fine-grain level when communities assemble from the same species pool, and at coarse-grain level when communities assemble from different species pools.

## Discussion

During the assembly of complex microbial consortia, selection will act on traits that enable or enhance growth on the available resources and/or on the secreted metabolic by-products (Fig. 1). By phenotyping a library of proteobacterial isolates across many single-nutrient environments we have shown that there exist similarities in growth rates (Fig. 2) and in by- product secretions when metabolically similar substrates are consumed (Fig. 3). By analyzing the composition of hundreds of experimentally assembled in-vitro communities, we have shown that quantitative similarities in metabolic traits lead to similarities in community composition with metabolically similar substrates selecting for taxonomically similar communities (Fig. 4). This result led us to propose and then validate a comparative approach for predicting community composition in new environments using communities assembled in metabolically similar environments (Fig. 5).

The space of all possible environments in which microbial communities may assemble is astronomically large and we have, by necessity, constrained our experiments to focus on a single type of variation; aerobic environments differing by a single carbon source. Despite this constraint, the comparative approach we have outlined holds promise due to the versatility of flux balance analysis. In principle, metabolic similarity can be computed for any arbitrary environment where the chemical composition is known and can be computationally defined. For example we were able to test our proposed comparative approach for environments containing 2-3 carbon sources and found that we are able to predict community composition on the nutrient mixtures from the composition observed in single nutrient environments (Fig. S21, Methods).

Future experimental work, assembling communities along alternative axes of environmental variation (such as different abiotic stressors) will be needed to determine whether similar degrees of predictability are observed along other axes of environmental variability.

Our results also suggest conditions for and limits on the predictability of community structure. In our experimental system, community assembly is predictable at coarse phylogenetic levels even when the pool of colonizing species comes from different inocula (Fig. 5). This predictability occurs due to phylogenetic conservatism of quantitative metabolic traits, which results in different strains from the same clade occupying the same functional role (Fig. 2B, Fig. S9) (Estrela, Vila, et al. 2022; Parras-Moltó and Aguirre de Cárcer 2021). If the quantitative traits influencing composition were more idiosyncratically distributed across the phylogeny, or across metabolically similar environments, then we would expect community composition to be far more unpredictable (Bittleston et al. 2020; Javdan et al. 2020). Interestingly, of the two major clades observed in our communities, P+ has much more variable by-product secretions than E+ (Fig. S9, Fig. 3E). We speculate that this may explain why communities in which P+ reached a higher abundance tended to be more variable at higher levels of taxonomy than communities in which E+ reached a higher abundance, despite this not being true at lower levels of taxonomic organization (Fig. S22).

In essence, our study establishes that the influence of different environments on microbiome composition are predictable when the traits determining composition are similarly distributed across quantitatively similar environments. Future empirical work is needed to elucidate whether these conditions hold true in more complex environments where a larger number of different traits are presumably under selection. We propose that high-throughput quantitative measurements of metabolic traits across diverse environments will play a critical role in identifying which environmental variables lead to idiosyncratic effects and which lead to predictable effects. Ultimately, our ability to predict and control microbiomes hinges on whether we can tame the immeasurable variability of complex environments. Encouragingly, our results imply that this challenge may not be insurmountable.

## Materials and Methods

### Seven carbon source community assembly experiment

For the seven carbon source community assembly experiment (Fig. 1), we followed the protocol outlined in Goldford et al 2018 which we recapitulate here. Communities were taken from 12 different soil inocula and serially transferred in M9 minimal media supplemented with one of seven different carbon sources (D-Glucose, Pyruvate, Citrate, L-Glutamine, Acetate, L-Leucine and L-Phenylalanine). All carbon sources were added at equal c-molar concentrations (0.07 C- mol/L). Communities were initialized by inoculating 4 µl of each source community into 500 µl of fresh minimal media. The communities were grown in 96 deep well plates (VWR) at 30°C without shaking for 48hr. At the end of the 48 hr growth cycle, 4 µl was transferred to 500µl of fresh media. This was repeated for a total of 12 transfers (∼84 generations). As in our previous work, for the first two transfers 200µl/ml of cycloheximide was added to eliminate eukaryotes. At the end of the experiment samples were stored by spinning down in a microcentrifuge for 10 min at 14,000 RPM at room temperature and storing cell pellets at -20°C. DNA extraction, library preparation and 16s rRNA gene sequencing was done as described previously (Goldford et al 2018).

Demultiplexed fastq files were processed using DADA2 version 1.6.0. The forward and reverse reads were truncated at position 240 and 160 respectively and merged with a minimum overlap of 100 bp (other parameters set to default values). Taxonomic assignment of each ESV was done using a naive Bayesian classifier method trained on the SILVA reference database (version 128). All samples were rarefied to the highest depth possible of 19969 reads.

### Strain Library and Phylogenetic Tree Construction

We started with a collection of 112 proteobacterial isolates, 77 of which had been isolated from glucose-enrichment communities and 35 of which were isolated directly from environmental samples (including soil, leaf litter, and pond water). The enrichment community isolates are the same as those studied in Estrela et al 2022. The soil isolates are new to this study and were obtained from 12 different environmental samples collected from around New Haven, CT (U.S.A.). Leaf, soil and river-bed samples were collected using sterile shovels and placed in 50 ml falcon tubes. Samples were then resuspended by vortexing in 50 ml of PBS. 2 µl of the sample suspension was then plated on DM1000 agar plates. From each sample, isolates with unique colony morphologies were picked after 48 hrs and streaked twice onto TSA plates to obtain pure cultures. For long term storages all strains were first grown for 48hrs on TSB and then stored at -80°C with 40% glycerol. Full-length 16S rRNA gene sequencing of every strain was conducted using the single colonies grown on TSB plates (GENEWIZ).

To ensure that our analysis was limited to strains which had high quality unique full-length 16s rRNA gene sequences, we developed a custom bioinformatics pipeline(Process_GENEWIZ.py). Firstly, low quality bases were removed from the start and end of the forward and reverse reads using the biopython implementation of the motts modified trimming algorithm with the default cutoff of 0.05 (Ewing and Green 1998). Next, forward and reverse reads were aligned using pairwise local alignment (pairwise2.align.localms) and merged. When merging sequences we used the most high confidence base and calculated the posterior probability for each base by combining the probability scores for the forward and reverse reads (Edgar and Flyvbjerg 2015). Taxonomy was assigned to these merged sequences in Dada2 using both the RDP dataset (training set 18) and the Silva 16S dataset (version 138). The two datasets gave the same taxonomic assignments at the Family level or higher, though we observed discrepancies at the Genus level. We choose to use the RDP assignment in the main text to ensure consistency with our previous work (Estrela et al 2022). None of our results depend on this choice.

In order to identify potentially duplicate sequences in the set of 112 merged sequences, we aligned every pair using the NeedleCommandline function in biopython and quantified the number of mismatches. We conservatively considered mismatches as a pair of non-identical bases which both had a phred quality score >10 (i.e a probability of being incorrect less than 0.05). Starting with these 112 sequences, we then iteratively removed sequences one by one in order to obtain a final set in which there was at least one high confidence mismatch between every pair. When choosing which sequence to remove from our analysis we removed shorter sequences and sequences for which phenotypic data had not been previously collected (Estrela et al 2022). We also removed one sequence that was substantially shorter than all other sequences (880 base pairs) even though it had no homologs. Removing these potential duplicate and lower quality sequences gave us 62 full-length 16s rRNA gene sequences which were between 1392 and 1472 base pairs in length. 49 of these strains are from Glucose enrichment communities and 13 are new soil isolates.

To construct a phylogenetic tree for this set of 62 strains, we performed multiple sequence alignment using clustalw version 2.0. We inferred a Maximum Likelihood phylogenetic tree using IQTREE2 version 2.1.2 with 1000 bootstrap replicates (iqtree -s alignment.fasta -bb 1000)(Minh et al. 2020). The tree was inferred under the TIM3+F+R4 model, which was selected as having the lowest BIC by ModelFinder(Kalyaanamoorthy et al. 2017). Bootstrap replication was performed using UFBoot and the consensus tree was constructed by averaging across the 1000 bootstrap replicates. The consensus tree was rooted using the 2 *betaproteobacteria* strains as the outgroup as the 60 other isolates are all *gammaproteobacteria.* All subsequent analysis was conducted using the rooted consensus tree.

### Growth Rate Quantification

Isolates were revived from glycerol stocks by streaking them onto chromogenic agar and growing them at 30°C for 24 hr (HiCrome Universal differential Medium from Sigma). Single colonies of each isolate were used to inoculate 500 μl LB (Lennox) in a 96 deep-well plate.

These pre-cultures were incubated at 30°C with shaking at 200 rpm for 24 hr. After 24 hr 100 μl of each sample was collected to measure Optical density (OD) 620 (which had reached between 0.2 and 1.5 across all cultures). Pre-Cultures were then diluted 1:10000 in M9 supplemented with one of the 19 different carbon sources. The final volume for the growth assays was 100 μl in 96 well plates. OD620 measurements were performed at 30 minute intervals over a 48 hr period with an Epoch 2 microplate spectrophotometer (BioTek). Between readings culture plates were stored with lids on in a Microplate Stacker (Bioplate) at 30°C without shaking.

On acetate as a substrate, two strains displayed large spikes in growth during the first couple of hours and so these growth curves were removed from the subsequent analysis. From the growth curve of every isolate (i) on each carbon source (ɑ) we estimated the average growth rate:

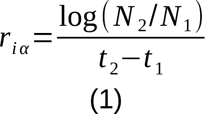

In this equation, t_2_ is the time at which we observed the maximum growth rate, N_2_ is the OD620 at this time point, N_1_ is the OD620 at the start of the growth curve (ignoring the first two measurements which eliminates artifacts from bubbles) and t_1_ is the corresponding time point. t_2_ was estimated by first by fitting a generalized additive model with an adaptive smoother to each growth curve using the *gam* function from the *mgcv* package in R and then taking the derivative of this smoothed curve to determine the time point at which the maximum derivative was detected. To avoid artifacts from measurement and fitting noise when strains are at low density, we set a minimum t_2_ to 5 hours.

Our reasoning for using this measure of growth rate is that it incorporates lag time differences for different strains (which would not be captured in the maximum growth rate) and it also captures differences in the time taken to consume different carbon source which would not be captured if we had used the average growth rate up to a fixed time point (such as the 16hrs used in (Estrela et al. 2021) . By ignoring timepoints after maximum growth rate has been achieved (t_2_) this measures also has the advantage of minimizing the confounding effects of growth on metabolic by-products (i.e diauxie) which would have been included in an area under the curve calculation of if we have fitted a logistic function to the entire growth curve .In Figure S7, we show that the relationship between metabolic similarity and growth rate similarities held when using these four alternative methods for quantifying growth. This includes measuring average growth rate at the fixed timed point of 16 hours (as in (Estrela et al. 2021); using the maximum growth rate estimated from the derivative of the smoothed curve (as in (Estrela, Vila, et al. 2022)); using the growth rate estimated by fitting a logistic function to each growth curve using the *growthcurver* package (Sprouffske and Wagner 2016); and measuring growth using the area under the curve estimated from *growthcurver* (Piccardi, Vessman, and Mitri 2019).

### Quantitative measure of metabolic similarity

For all 48 carbon sources included in this study, we used the cobrapy implementation of Flux Balance Analysis to obtain a predicted vector of fluxes through a microbial metabolic network (Ebrahim et al. 2013). For this analysis we started with the *E. coli* metabolic model iJO1366, a well benchmarked reference model of *E. coli (Orth et al. 2011)* that we had previously used to predict community assembly on D-Glucose (Estrela, Vila, et al. 2022). All simulations were conducted using the cobrapy package (Ebrahim et al. 2013). In order to obtain a flux vector for every carbon source, we supplemented this model with reactions from a universal model of microbial metabolism (Machado et al. 2018) (Table S1). For the majority of carbon sources the addition of a transporter and exchange reaction was sufficient to ensure growth. In a limited number cases, additional internal metabolic reactions had to be added and those metabolic reactions are listed in Table S1.

All carbon sources were supplied to the model at equal carbon concentrations by setting the lower bound on its exchange flux to -1 cmol/gDWh. Trace elements, oxygen, and nitrogen where supplied to the model in excess by setting the exchanges on the following metabolites to be unbounded: ca2_e, cbl_1 e, cl_e, co2_e, cobalt2_e, cu2_e, fe2_e, fe3_e, h_e, h2o_e, k_e, mg2_e, mn2_e, mobd_e, na1_e, nh4_e, ni2_e, pi_e, sel_e, slnt_e, so4_e, tungs_e, zn2_e, and o2_e. We set the lower bound on ATPM maintenance to 0 as we wanted to estimate the flux distribution when resources were in excess and so the effects of growth independent maintenance would be negligible. Growth was optimized using parsimonious flux balance analysis (cobra.flux_analysis.pfba), with the biomass reaction (BIOMASS_Ec_iJO1366_core_53p95M) set as the objective function.

From the predicted fluxes, we removed all transporters, exchange reactions and the biomass reaction in order to obtain a vector of intracellular metabolic fluxes. We calculate the metabolic similarity between every pair of carbon sources as the Pearson correlation in optimal intracellular metabolic flux distributions.

To determine how sensitive this quantitative measure of substrate similarity is to the specific model organism used as a reference, we repeated the above analysis with 4 different unmodified high quality metabolic models downloaded from the BiGG database. These models were iML1515 (*E. coli*), iYL1228 (*K. pneumoniae*), STM_v1_0 (*S. enterica)* and iJN1463 (*P. putida).* We conducted flux balance analysis for the 19 carbon sources shown in Figure 2.

These models varied in the number of carbon sources on which they could grow (13/19 - 17/19). For carbon sources on which the model could grow, metabolic similarities were highly correlated with the metabolic similarities calculated using the modified model (which could grow on all carbon sources We therefore use the similarities calculated with the modified model for all of our analysis in the paper.

### Glucose byproduct quantification

The glucose by-products quantification (Fig. S9) combines publicly available data (Estrela et al 2021) as well as new measurements for the new isolates collected for this paper. As previously described, isolates were all preconditioned to grow on glucose minimal media (500 μl) for 48 hr. Then, 4 μl of the preconditioned cultures was inoculated into 500 μl fresh glucose media. 100 µl samples were collected after 16 hr and the OD620 was measured. The remaining sample was centrifuged at 3000 rpm for 25 min to separate cells from supernatant. Supernatants were transferred to two 96 well plate 0.2 µm AcroPrep filter plates on top of a 96 well NUNC plate fitted with the metal collar adaptor and centrifuged at 3000 rpm for 10 min. The supernatant was immediately frozen at -80°C until processing. For the 13 new natural isolates glucose, acetate, lactate and succinate concentrations were repeated as described previously (Estrela et al., 2022). Gluconate concentrations for all strains are new to this paper and were measured using a D-Gluconate assay kit (ab204703).

2-Ketogluconate concentrations were measured using a protocol adapted from Molina et al. (2019). Briefly summarized, 50 μl of o-phenylenediamine dihydrochloride was combined with 100 μl of diluted filtered culture supernatant and heated at 100°C for 30 minutes. The absorbance of the reaction mixture was measured at 330 mm and 2-ketogluconate concentrations were estimated by comparing the diluted supernatant to a standard curve.

Interference from other metabolites (D-Glucose and D-Gluconate) was detected by quantifying the ratio of OD330 and OD360 and removing data points where the ratio was greater than 2 (Lanning and Cohen 1951).

### Targeted Metabolomics

We revived *E. coli* MG1655, *P. putida* KT2440, an *Enterobacter* sp. and a *Pseudomonas* sp. (two isolates from the glucose communities in Goldford et al. [2018]) on LB Agar. For each strain we picked two replicate colonies and inoculated them into 50 ml falcon tubes containing 5 ml of LB (Lennox). Falcon tubes were incubated at 30°C (shaking) for 16 hrs. After this all 8 populations were brought into balanced exponential phase by diluting (1:5) three times into fresh LB. The first three dilutions were performed at 1 hr intervals after which cultures were allowed to grow for an 1hr and 30min. At this point, cells were centrifuged and washed three times in M9 minimal media containing no carbon source to remove any leftover LB before being resuspended in M9 Minimal media. Cell densities were normalized to a pre-inoculation OD of 0.1.

All eight samples (four taxa, two replicates) were used to inoculate 3 replicates each, which were grown at 30°C in 500 μl of M9 media containing one of five possible carbon sources (D-Glucose, D-Fructose, Glycerol, Pyruvate and L-Malate). Each replicate was used for a different time point (16 hr, 28 hr and 48 hr). At each timepoint, OD readings were taken and the supernatant was extracted as described previously. Spent media was submitted for a targeted metabolomics analysis carried out by the Metabolomics Innovation Center (TMIC), in Alberta, Canada. Metabolite quantification in samples was conducted using liquid-chromatography mass spectrometry (LC-MS) as described in Estrela et al. (2022). Measurements were also taken on D-Arabinose and D-Ribose for the *Enterobacter* and *E. coli* MG1655 after 28 and 48 hrs of growth. Measurements were also taken on Succinate and Fumarate for the *Pseudomonas* and *P. putida* KT2440 after 28 and 48 hrs of growth.

### Untargeted Metabolomics

A *Klebsiella* and a *Pseudomonas* isolate from Estrela et al. (2022) were revived on chromogenic agar. Three replicates of each isolate were preconditioned for 48hr on minimal media containing one of the following 19 carbon sources: D-Glucose, D-Fructose, D-Galactose, D-Ribose, L-Arabinose, Glycerol, Gluconate, 2-Ketogluconate, Pyruvate, D-Lactate, Citrate, Fumarate, L-Malate, Acetate, L-Glutamine, L-Leucine, or L-Phenylalanine as well as a no-carbon source control. After preconditioning, 4 μl of culture was transferred to fresh media and grown for 24 hr before spent media extraction. Growth conditions and spent media extraction were identical to the previous assays.

Samples on which significant growth had been observed (OD620>0.04) were subjected to untargeted metabolomics (LC-MS). The analysis was run on an Agilent 6550 Q-TOF mass spectrometer operated in negative mode. Mass analysis was acquired from 50-1100 m/z, with a gas temperature of 275°C, gas flow of 11 L/min, nebulizer pressure of 30 psig, sheath gas temperature of 350°C, and sheath gas flow of 12 L/min. Capillary voltage was 3500 V, nozzle voltage was 2000 V, with the fragmentor set at 365 V. Spectra were corrected using the Agilent API-TOF reference mass solution kit. The LC stack consisted of a 1290 Binary pump, 1290 autosampler thermostatted at 8 degrees C, and a column heater set at 35°C. The column was a Phenomenex Kinetex F5 2.1 mm * 100 mm column. The LC flow was 0.2 mL/min, with mobile phase A is 0.1% formic acid in water, and mobile phase B is 0.1% formic acid in acetonitrile.

The gradient is as follow:

Hold at 100% A from 0 minutes to 2.1 minutes

Ramp from 100% A to 5% A from 2.1 minutes to 14 minutes Hold at 5:95 A:B from 14 minutes to 16 minutes

Return to 100% A from 16.0 minutes to 16.1 minutes Hold at 100% A from 16.1 minutes to 20 minutes

Data was converted to .mzml using MSConvert software version 3.0.21071-c909413d7, and metabolites were integrated using El-Maven software, version 0.12.0.

Across all samples peaks mapping to 69 different metabolites were detected, annotated and integrated. 14 of these peaks were found at significant abundances in the no carbon source controls (>5% maximum peak height across all samples), so we removed these from the subsequent analysis. Samples from the *Enterobacter* and the *Pseudomonas* were always analyzed separately. For the analysis shown in Figures 3B-E and Figure S11 peak heights were averaged across all 3 replicates on each carbon source and then normalized across carbon sources (Z-Score Normalization). For Figures S12 and S13 the normalization was conducted across all samples (i.e including all 3 replicates separately..

### 42 Carbon Source experiment

Three communities each taken from different soil inocula were first grown in Tryptic Soy Broth (TSB) for 24 hr. Each community was then used as inoculum for M9 minimal media supplemented with equal c-molar concentrations (0.07 C-mol/L) of one of 42 different possible carbon sources .These carbon sources were: Benzoate, Methanol, Ethanol, 1- Propanol, Butanol, Glycolate, Galactitol, Propionate, Acetate, Formate, L-tartrate, Hexanoate, Decanoate, Ribitol, Myo-Inositol, D-Mannitol, Pyruvate, Melibiose, D-Glucose, D-Fructose, D-Galactose, L- Lactate, D-Sorbitol, Salicin, Cellobiose, D-Arabinose, L-Arabinose, Lactose, 2-Oxoglutarate, Trehalose, Sucrose, Glycerol, Raffinose, L-Rhamnose, D-Ribose, Maltose, Citrate, Fumarate, Succinate, L-Malate, Butyrate and Glyoxylate. We note that several communities were also assembled on Tween-80 and Isopropanol but we removed these communities from our analysis because these metabolites are currently absent in the BIGGs database and so a metabolic similarity measure could not be computed.

The communities were grown in 96 deep well plates (VWR) at 30°C without shaking for 48 hr. At the end of the 48 hr growth cycle, 4 µl was transferred to 500 µl of fresh media. This was repeated for a total of 10 transfers (∼70 generations). For the first two transfers 200µl/ml of cycloheximide was added to eliminate eukaryotes At the end of the experiment, samples were stored by spinning down in a microcentrifuge for 10 min at 14,000 RPM at room temperature and storing cell pellets at -20°C. DNA extraction, library preparation and 16s rRNA gene sequencing was done as described in the previous section (Seven carbon source community assembly experiment).

One of the 126 samples had an extremely low read-depth (786 reads) and was discarded. In order to keep samples from as many different carbon sources as possible we choose to rarefy our samples to the highest depth possible of 2733 reads. We could have used a higher read-depth, but this would have required discarding additional samples which would have reduced the statistical power of our analysis. Furthermore, because we measure community similarity using Renkonen similarity, further increasing read-depth would not significantly improve our resolution as Renkonen similarity is insensitive to rare taxa.

### 24 Carbon Source experiment and mixed nutrients experiment

This data came from enrichment community experiments that we have previously described and reported elsewhere (Estrela, Diaz-Colunga, et al. 2022) Following the same procedure described for the seven carbon source community experiment, eight soil samples were used as inocula for enrichment community experiments in one of 24 different carbon sources which were diluted every 48 hrs for 10 transfers (∼70 generations) in four replicates. To allow for a comparison with the other experiments, the analysis shown in Figures S18-S20 uses one of the four replicates in this dataset (replicate 1) but our results are quantitatively similar on all four biological replicates.

The 24 carbon sources used in this experiment were: D-Glucose, D-Mannitol, Cellobiose, Trehalose, D-Fructose, D-Ribose, L-Arabinose, D-Arabinose, Salicin, Glycerol, Galactitol, Myo-Inositol, Citrate, 2-Oxoglutarate, Succinate, Fumarate, Acetate, Propionate, Benzoate, L-Glutamine, L-Valine, Glycine, L-Tryptophan and L-Malate.

Two of the eight inocula were additionally used for enrichment experiments containing mixtures of two or three different carbon sources(Estrela et al. 2021; Estrela, Sánchez, and Rebolleda-Gómez 2021). The total carbon content in this environment was the same as the single nutrient environments (0.07 C-mol/L). Each carbon source in this experiment was mixed at equal c-mol concentration (i.e 0.035 C-mol/L in the two carbon sources and 0.023 C-mol/L in the three carbon source experiment). The two carbon sources treatments contained either D-Glucose or Succinate mixed with one of eight different carbon source: D-Cellobiose, D-Fructose, D-Ribose, Glycerol, Fumarate, Benzoate, L-Glutamine, and Glycine. The three carbon source treatments contained D-Glucose and Succinate mixed with one of the same eight carbon sources.

DNA extraction and sequencing for all these communities after 10 transfers was conducted as described in our previous work (Estrela et al. 2021; Estrela, Diaz-Colunga, et al. 2022).

Demultiplexed fastq files were processed using DADA2 version 1.6.0 .The forward and reverse reads were truncated at position 240 and 160 respectively and merged with a minimum overlap of 100 bp (other parameters set to default values). Taxonomic assignment of each ESV was done using a naive Bayesian classifier method trained on the Silva 16S reference database (version 128). All samples were rarefied to the highest depth possible of 10779 reads.

### Statistical analysis of correlations

All statistical analysis was conducted in R Version 4.3.1. Correlations were calculated using the base R *cor* function and used all complete observation pairs (use= “pairwise”). The statistical significance of correlations between difference matrices (i.e Figure 2F, 3 was determined using the mantel permutation test (*mantel.rtest* in the *ade4* package) with 9999 permutations (*nrep =9999*) (Mantel 1967).

### Measure of community similarity

The similarity between community pairs was measured by calculating Renkonen similarity at different taxonomic scales. This is defined as:

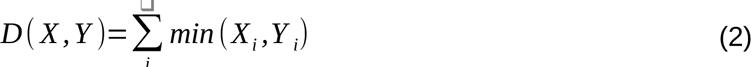

Where X_i_ and Y_i_ are the relative abundances of Taxon i in communities X and Y respectively. Because this is mathematically equivalent to 1 - the Bray-Curtis dissimilarity we calculated Renkonen similarity using the *vegdist* function in the R vegan package (Oksanen 2010).

Throughout the paper we used the one-tailed Mann-Whitney U-Test (R Function w*ilcox.test, exact=FALSE,* alternative= ‘greater’) to determine if the pairwise similarities for one set of communities (i.e same inoculum) were significantly larger than for another set of communities (i.e different inocula). The same statistical test was used to compare the observed distribution of performances to the null distribution of performances in Figures 5, S19 and S20.

### Decomposing the variance in community composition

The proportion of variation in community composition (beta-diversity) explained by the inoculum and carbon source were calculated at different taxonomic resolutions using the *varpart* function in the R vegan package (Oksanen 2010). *Varpart* partitions the variation in community data using the adjusted R^2^ calculated from canonical redundancy analysis (Borcard, Legendre, and Drapeau 1992; Legendre 2008).

### Predictions of community composition

The ability to predict community composition using a comparative approach was assessed across pairs of inocula using “leave-one out carbon-source out cross-validation”. Let X be the set of target carbon-sources on which inoculum A has been assembled and Y be the set of template carbon-sources on which inoculum B has been assembled. For each target carbon-sources in X, we find the most metabolically similar non-identical template carbon-sources in Y. The composition of the community assembled from inoculum A in the target environment is then predicted to be the same as the composition of the community assembled from inoculum B in the template carbon-sources. This approach is equivalent to using a nearest neighbor classifier with metabolic similarity as the measure of distance in the feature space (metabolic flux space).

For each inoculum pair, the performance of our predictions was quantified by measuring the Renkonen similarity between predicted and observed community composition for all target environments in X. We use this measure of performance to account for the compositional nature of the data that is being predicted (i.e relative abundances). As a null model we compared this observed distribution to the distribution of Renkonen similarity between all pairs of non-identical carbon-sources across X and Y. This null is analogous to repeating the procedure for making predictions except community composition for each target carbon-source in X is predicted using a randomly chosen non-identical template carbon-source in Y irrespective of metabolic similarity.

In Figure S20 we show predicted vs observed composition on 18 different mixed carbon source environments made using all the single carbon source environments assembled from the same inoculum (B) or different inoculum (C). Metabolic fluxes for the target environments (the mixtures) were quantified by setting the lower bound on the exchange flux to -1 cmol/gDWh for the nutrient pairs and -0.033 cmol/gDWh for the nutrient triplets. This reflects the fact that total carbon concentration was kept constant across environments. The template single environment was chosen by identifying the most metabolically similar single nutrient environment to the mixed nutrient environment. For these predictions, carbon sources included in each of the mixture were not used as templates. For example, for the D-Glucose and D-Ribose, mixture composition was predicted using the 22 single-nutrient environments excluding D-Glucose and D-Ribose. The null distribution was determined using the same procedure as the single nutrient environment.

## Acknowledgements

We thank all members of the Sanchez, Mehta, Petrov and Segre lab for helpful discussion. We thank Paul Turner, David Post, Gunter Wagner, Steven Stearns as well other members of the Ecology and Evolutionary Biology Department at Yale for useful feedback. We thank Christopher Singleton and The Metabolomics Innovation center for providing untargeted and targeted metabolomic services respectively. We thank Juan Nogales for assistance with the 2-ketogluconate quantification and Daniel Machado for assistance with using the *CARVEME* package.

## Author contributions

JCCV, JG, SE, DB, RM, PM and AS conceived of the study. All authors helped design the experiments. JCCV, JG, SE, DB, ASG, NL,and MRG performed the experiments. JCCV, JG and SE analyzed the data with tools contributed by DB, ADS and RM. JCCV wrote the paper with input from AS. All authors discussed the results and reviewed the paper.

## Data and Code availability

All data and code needed to reproduce our results will be made available before publication at https://github.com/jccvila/Vilaetal2023. A Zenodo repository will be uploaded as of the date of publication. New raw 16S rRNA amplicon sequences and metadata files will be deposited on the NCBI SRA database and will be made publicly available as of the date of publication. Data shown in Figure S1 and Figure S9 has already been published (Estrela, Vila, et al. 2022). Data for the 24 carbon sources from the 8 different inocula and the carbon source pair from two different inocula has either already been published or will be deposited upon publication of a preprint currently under review (Estrela et al. 2021; Estrela, Vila, et al. 2022; Estrela, Diaz-Colunga, et al. 2022)

