## Supplementary Material for "Metabolic similarity and the predictability of microbial community assembly"

**Supplementary Materials for: metabolic similarity and the predictability of microbial community assembly**

This File includes:

- Supplementary Figures 1-22
- Supplementary Table 1

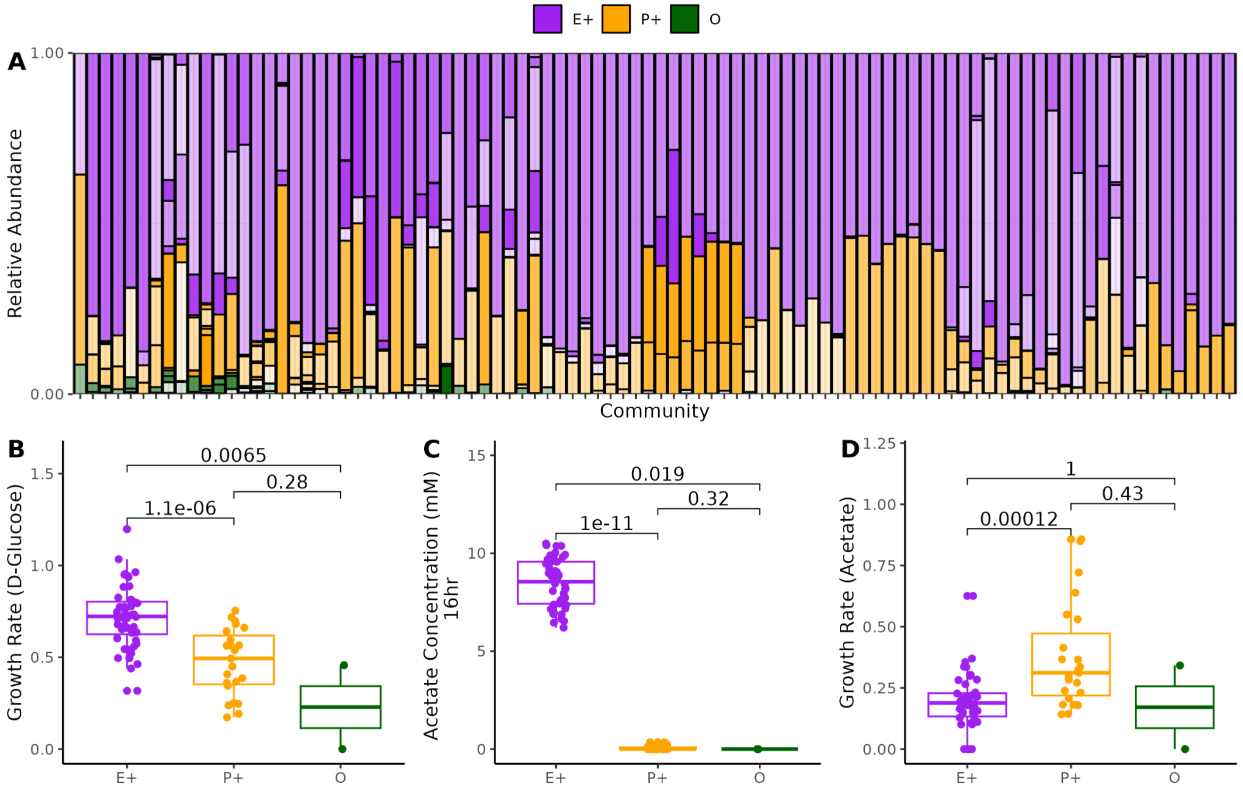


**Supplementary Figure 1: Community composition on D-Glucose depends on the taxonomic distribution of both growth rates and secreted metabolic by-products. (A)** Composition of communities assembled on minimal glucose is convergent at coarse phylogenetic levels, despite variability at lower taxonomic scales. Barplot shows the composition of the 92 enrichment communities from 12 different inocula after 12 growth/dilution cycles on minimal glucose. Purple corresponds to members of the 'E+' clade which includes Enterobacteriaceae and the closely related Yersiniaceae, Erwiniaceae and Aeromonadacae. Orange corresponds to members of the 'P+' which includes Pseudomonadaceae and the closely related Moraxellaceae. Green corresponds to the two members of a Betaproteobacteria clade. Different shades of the same color show different ESV within the same phylogroup. (**B–D)** We have previously shown that taxonomic convergence at this coarse-grain phylogenetic level can be explained via a metabolic self-organization where members of the 'E+' clade are selected for the ability to grow rapidly on glucose (Panel B) whilst secreting quantitatively similar levels of the organic acid acetate. (Panel C) . Members of the 'P+' clade are selected for the ability to grow rapidly on organic acids including acetate (Panel D). These figures are replotted using data from Estrela et al 2022.

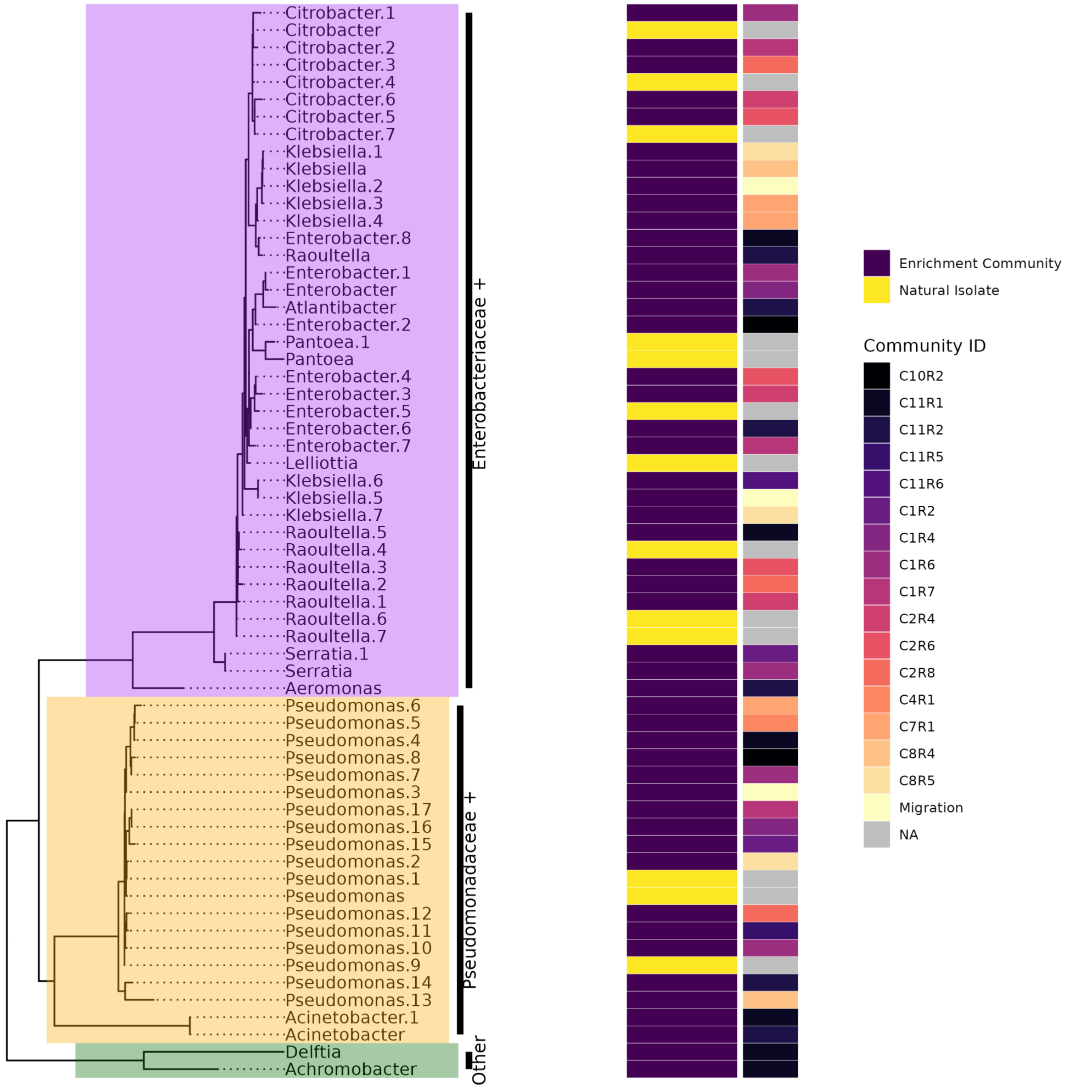


**Supplementary Figure 2: 16s Phylogenetic Tree of isolate library.** The strains used in this study were selected from a broader pool of 111 strains for which full length 16s length sequencing had been obtained. (Methods). The majority of the 62 strains were isolated from glucose enrichment communities as described in Estrela et al 2022. We also included 13 natural isolates that we sampled directly from different soil sources (Methods). A Maximum Likelihood phylogenetic tree was constructed using the full-length 16S sequence of each isolate (Methods). Heatmap shows the community of origin of each isolate. The community ID corresponds to the ID used in previously published work (Goldford et al. 2018; Estrela et al. 2022; Chang et al. 2023)

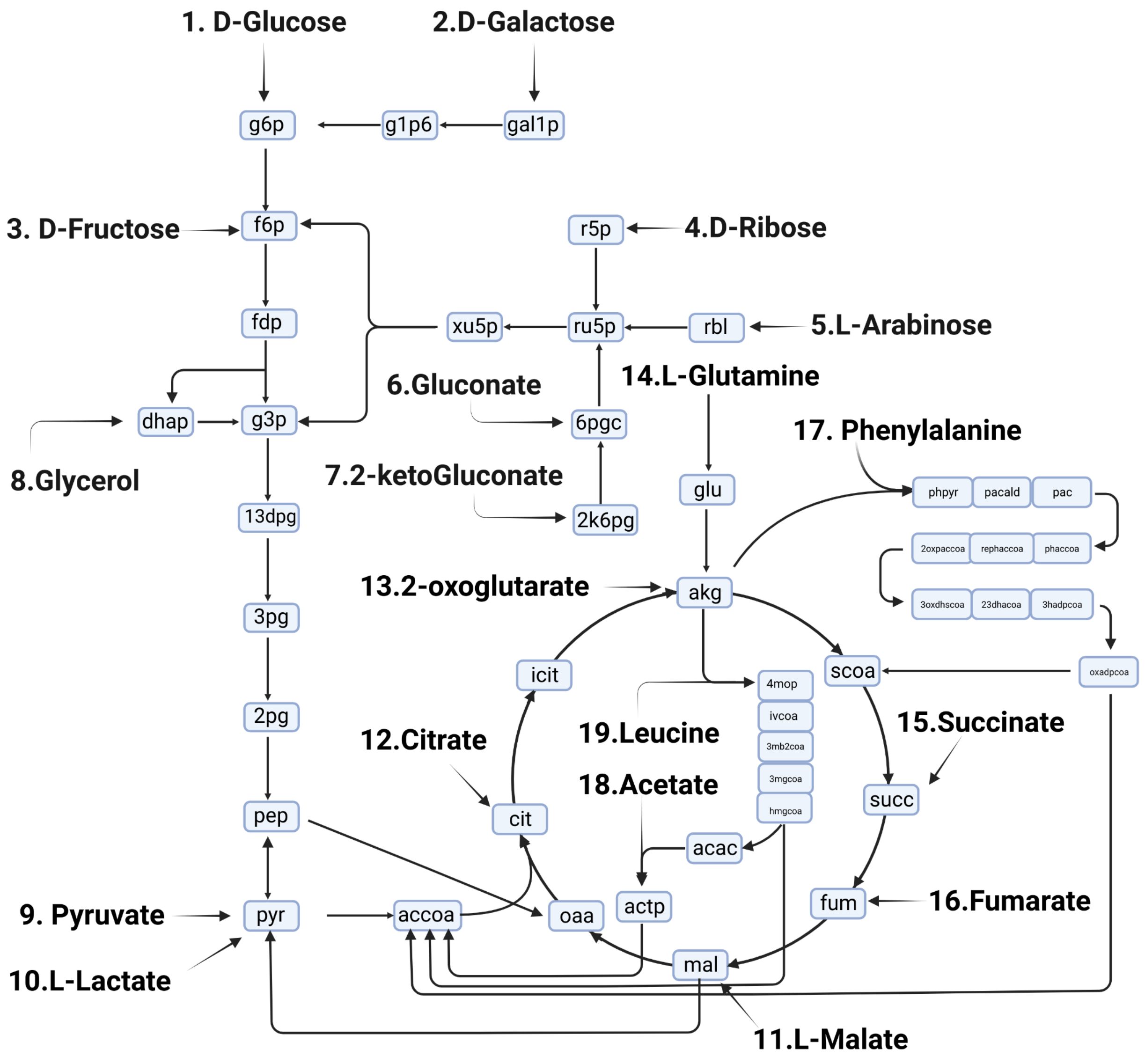


**Supplementary Figure 3: Metabolic network illustrating the metabolism of the 19 carbon sources used for the experiments in Figure 2 and Figure 3**. Carbon sources are numbered by the order of entry into glycolysis and the TCA cycle starting with D-Glucose. L-Malate is set as the first carbon source in the TCA cycle because it can be reversibly converted to Pyruvate by malate dehydrogenase (Sauer and Eikmanns 2005). We note that any ordering based solely on network topology will ultimately be arbitrary because metabolic networks are not tree-like. This further justifies the quantitative approach based on optimality that we have to use to define carbon source similarity in this paper (Figure 2C). Metabolites in this figure (nodes) are labeled using the ID from the BiGG database (King et al 2016).

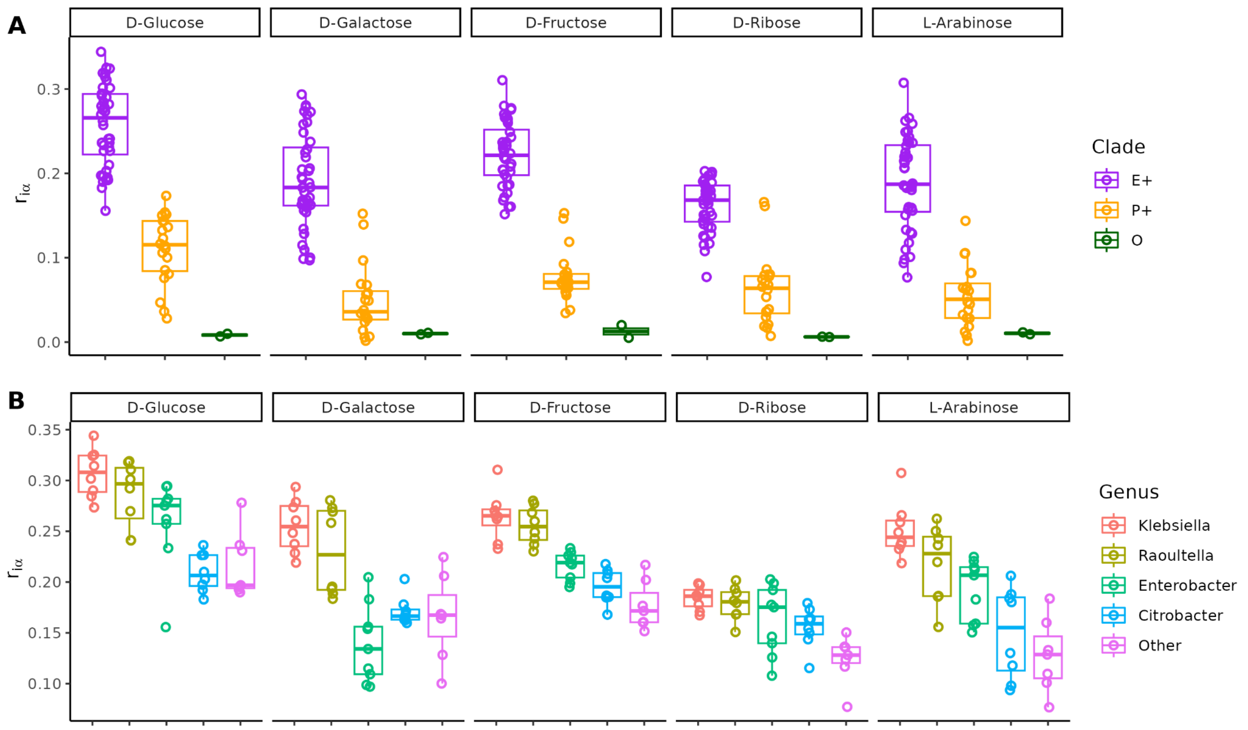


**Supplementary Figure 4: Taxonomic variation in the average growth rate on the five sugars. (A)** Members of the E+ Clade have a larger average growth rate on all 5 sugars than the members of the other two clades. **(B)** Within the E+ clade, members of the *Klebsiella* and *Raoultella* genus grow faster on all 5 sugars.

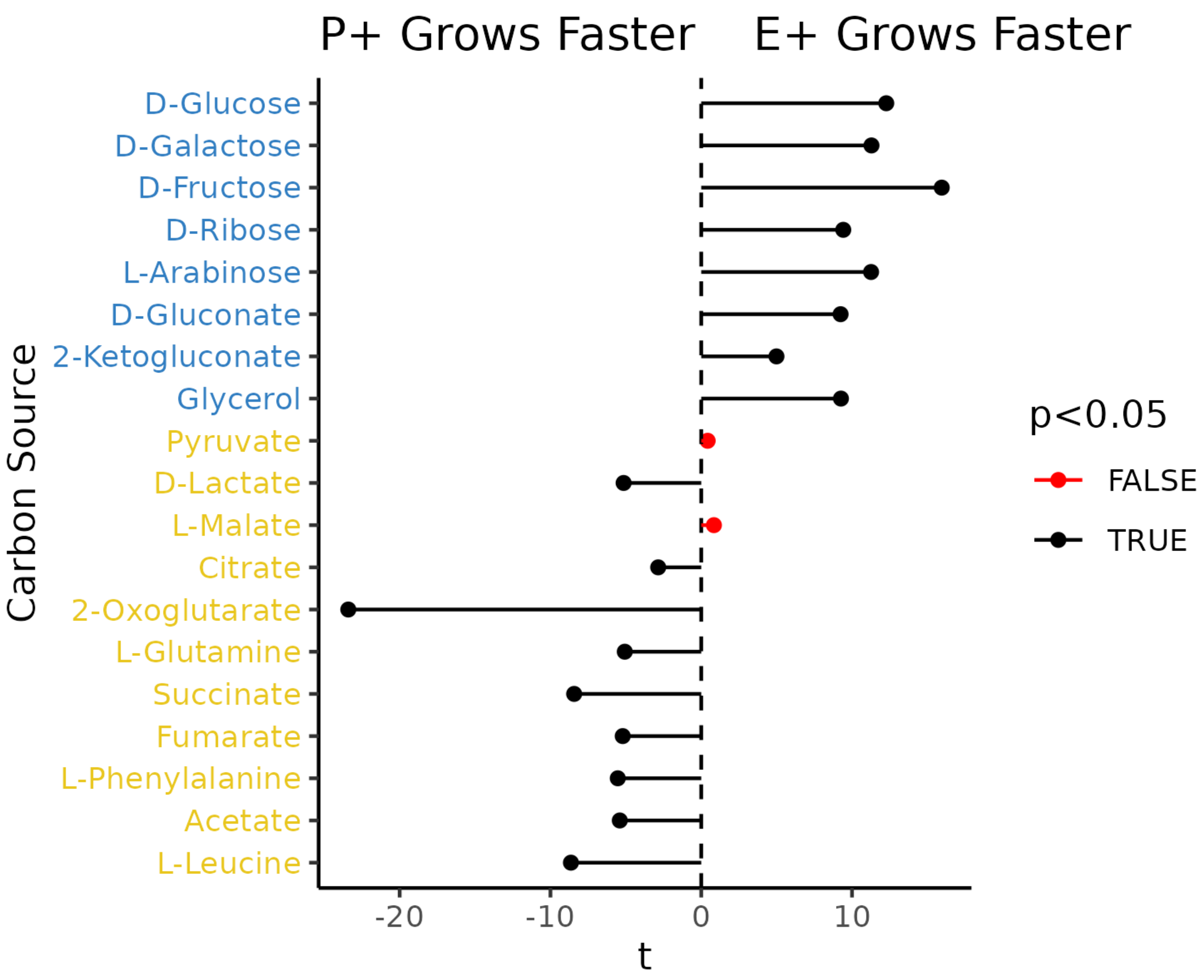


**Supplementary Figure 5: Test statistic (t) for Welch two sample t-test comparing growth rates of members of E+ vs P+ Clade.** Members of the E+ Clade grow significantly faster on the 8 Glycolytic Carbon sources (Blue). Members of the P+ Clade grow significantly faster on 9/11 Gluconeogenic carbon sources (Yellow). No significant differences are observed on Pyruvate or L-Malate. Carbon sources are ordered as in Fig S3.

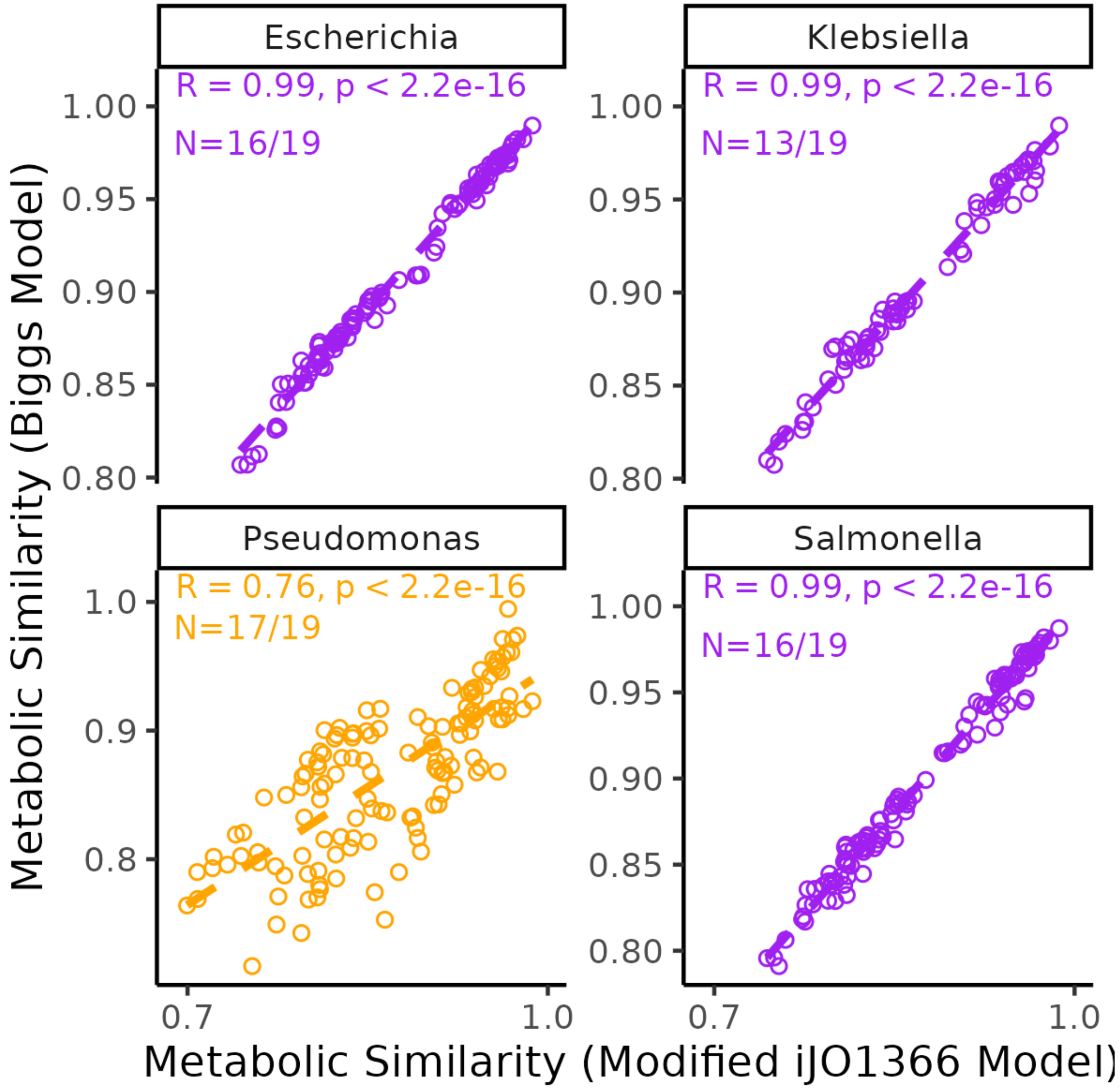


**Supplementary Figure 6: Substrate similarities are conserved across metabolic models from different organisms.** To quantify substrate metabolic similarity we performed flux balance analysis using the iJO1336 metabolic model for E.coli which we had modified to ensure that it could predict the growth on every carbon source used in this study. Metabolic similarity is defined as the correlation in predicted intracellular metabolic fluxes for the optimal growth on each carbon source. On the X axis we plot the predicted metabolic similarity for every pair of carbon sources calculated using this modified model. On the Y axis we plot the metabolic similarity for a pair of carbon sources calculated using an alternative (unmodified) metabolic model for different bacterial genera found in our communities. N gives the number of the 19 carbon sources studied in Figure 2 and 3 on which the model grew. Metabolic models were downloaded from the biggs database and correspond to *e.coli* (iML1515), *k.pneumoniae (*iYL1228), *s.enterica (*STM_v1_0) and *p.putida (iJN1463) (King et al. 2016).*

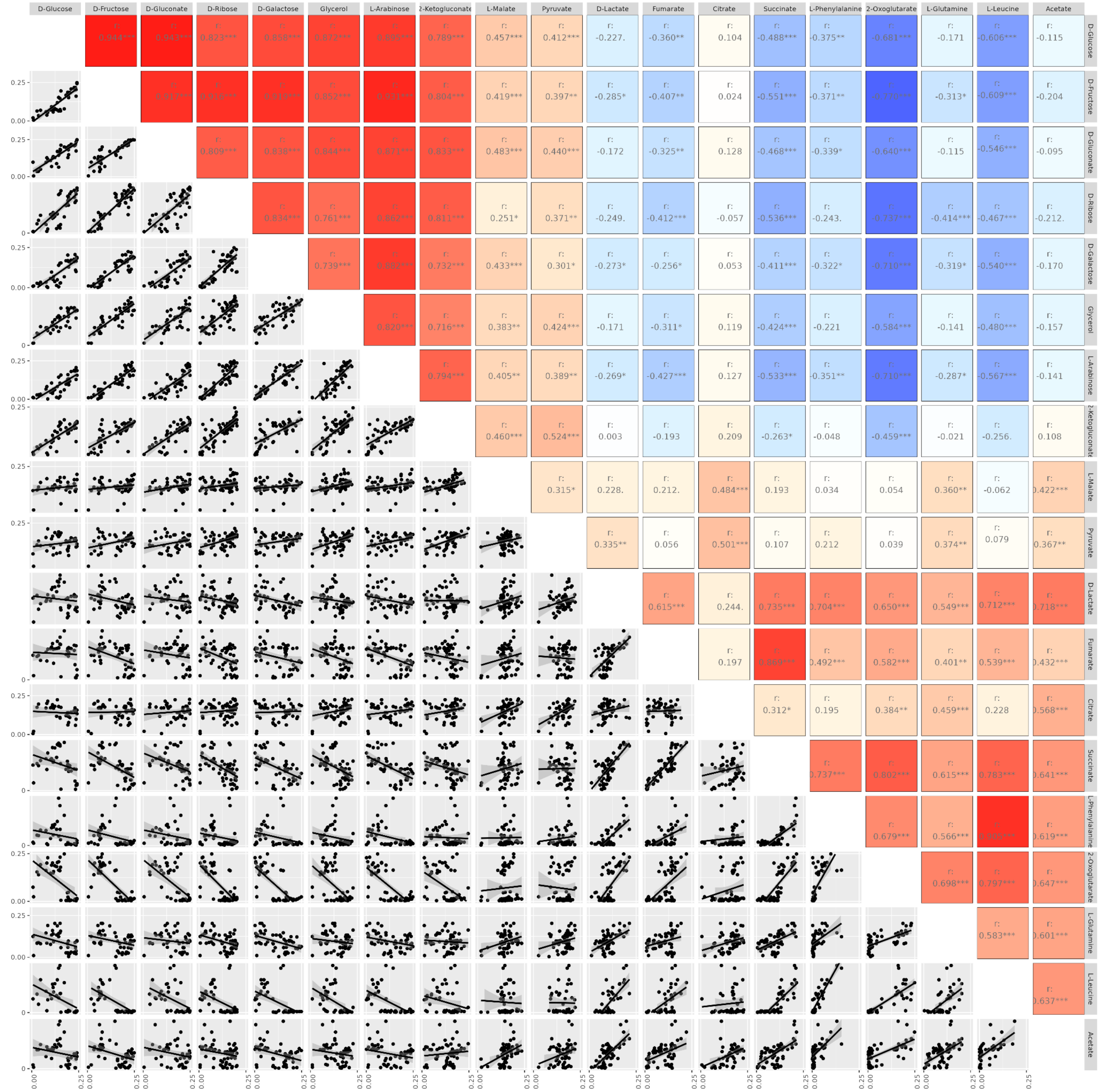


**Supplementary Figure 7: Correlation in isolate growth rate for all 171 carbon source pairs.** Bottom left: dot plot showing correlation in strain-specific growth rates for every pair of carbon sources, top right shows the direction of the correlation and whether it is significant for each carbon source (blue = more negative correlation, red = more positive correlation). In figure 2E we plot this relationship for D-Glucose and D-Fructose (top left panel).

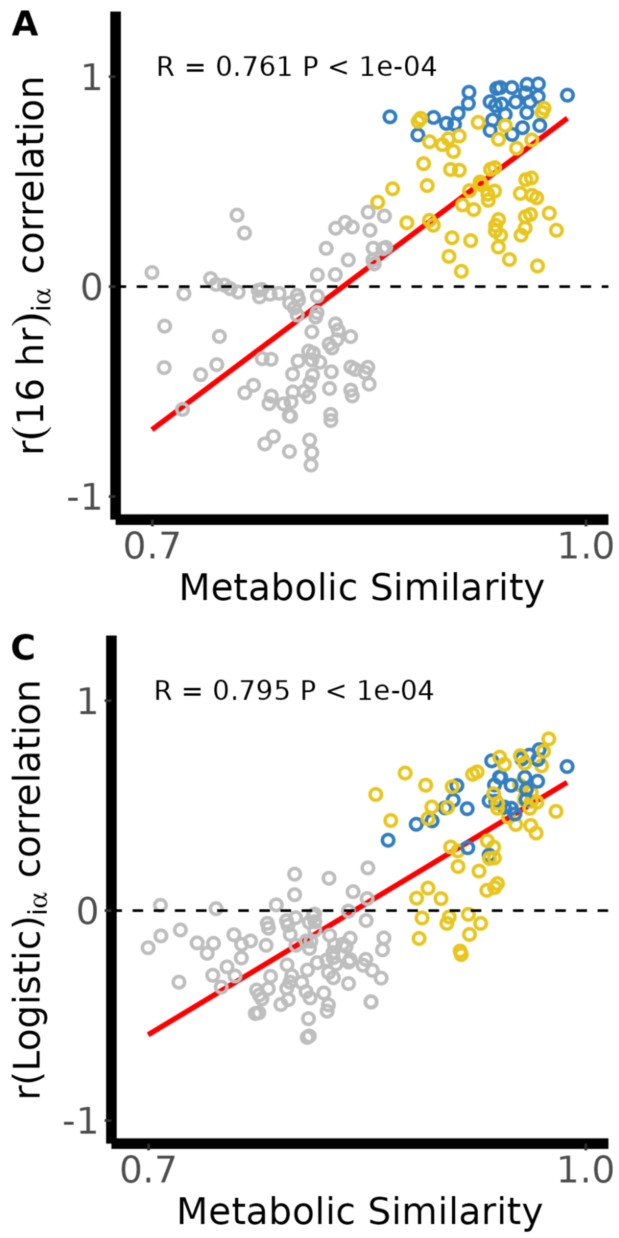


**Supplementary Figure 8: The correlation between metabolic similarity and growth ability is robust to alternative measures of growth capabilities.** Replotting of figure 2F with growth capability estimated using four alternative measures (Methods). A) Average growth rate up to 16 hrs; B) maximum growth rate estimated from the first derivative of the growth curve; C) growth rate estimated by fitting a logistic model; and D) Area under the curve. We always observed a positive correlation between metabolic similarity and the correlation in growth capabilities (Mantel test $p<0.001$).

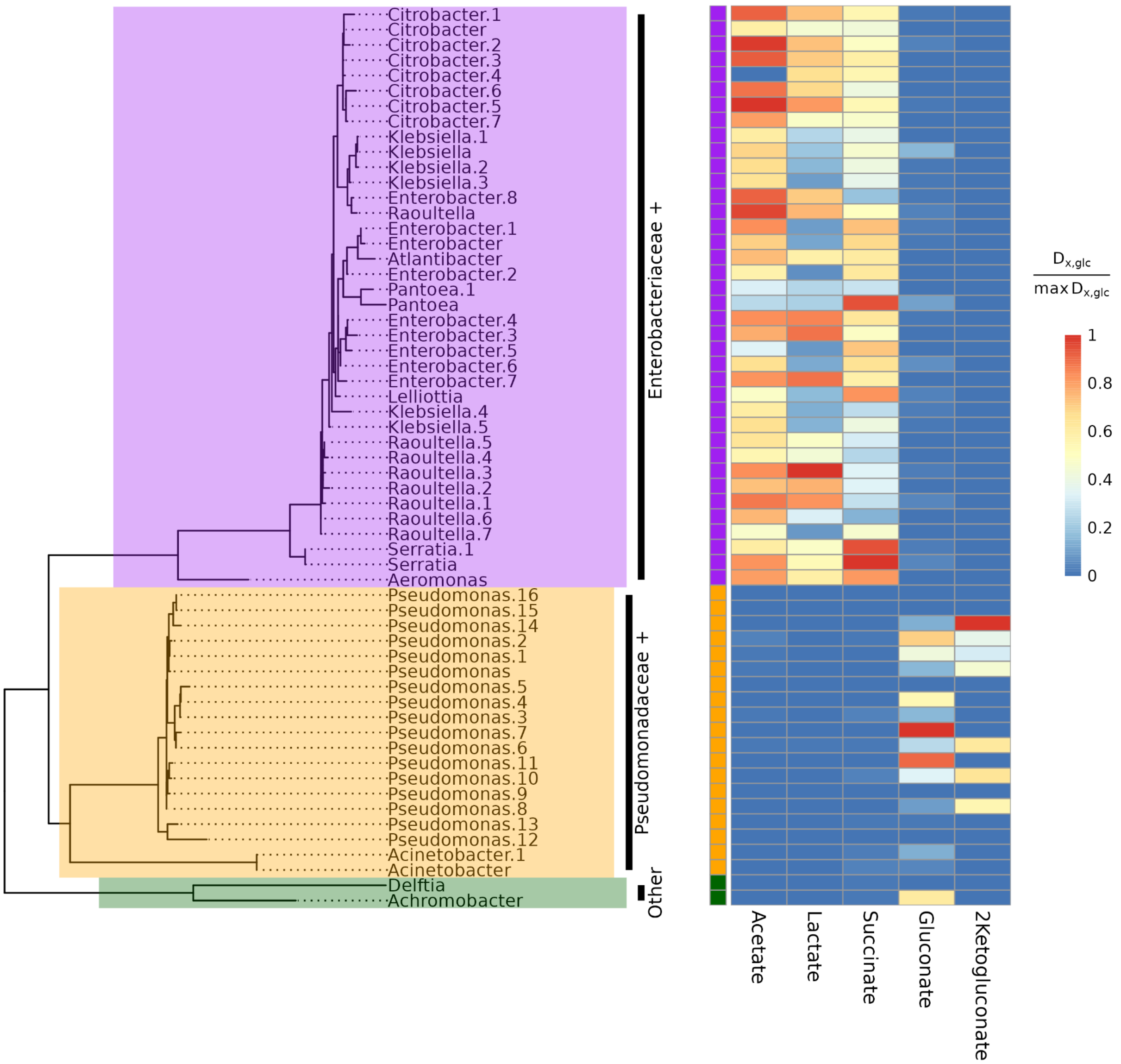


**Supplementary Figure 9: Metabolic secretions on D-Glucose differ between the two major clades.** Phylogenetic heatmap showing levels of acetate, lactate, succinate, gluconate and 2-ketogluconate secreted by 59 out of the 62 strains after 16hrs of growth on minimal glucose (Methods). E+ strains secrete higher level of Acetate(t=24.8,P <2.2e-16), Lactate (t=11.3,P=6.7e-14) and Succinate (t=8.7,P= 6.7e-12) whereas P+ strains secrete higher levels of Gluconate (t=-14,two sample T-test, one-tailed P =1.71e-12) and 2-ketogluconate (t=-2,P = 0.006). Reported P-values correspond to a two sample T-test (one-tailed). In order to better visualize the difference across the two clades the amount of each by-product per glucose consumed (in units carbon) is normalized to the maximum value across all strains.

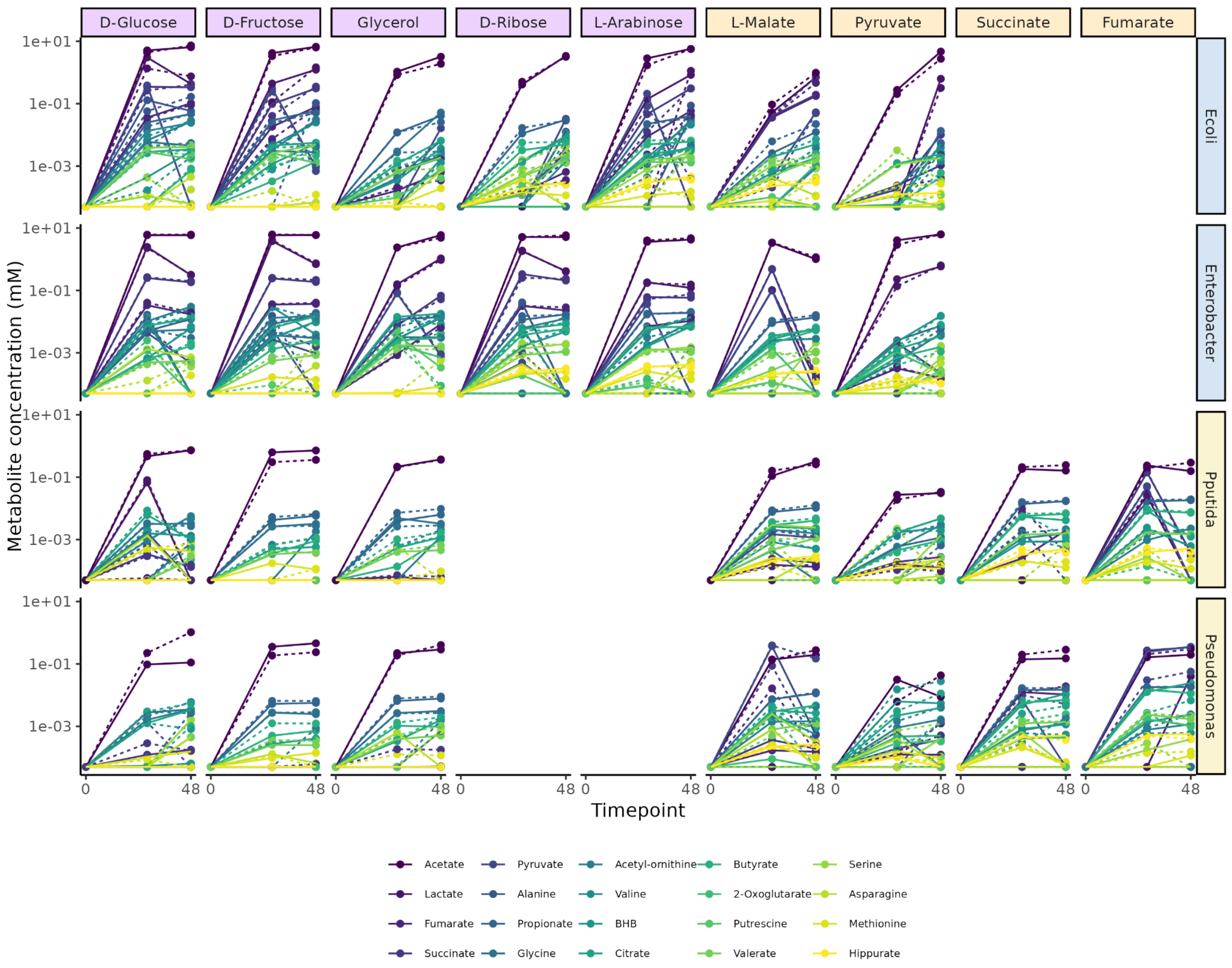


**Supplementary Figure 10:** Metabolite abundance estimated through targeted LC-MS for 4 strains on different carbon sources. Dotted and solid lines correspond to one of two different biological replicates. In Figure 3B we show the results for the Enterobacter on the 5 carbon sources (D-Glucose, D-Fructose, Glycerol, L-Malate and Pyruvate) for which we collected data at all 3 time-points and for all srains .

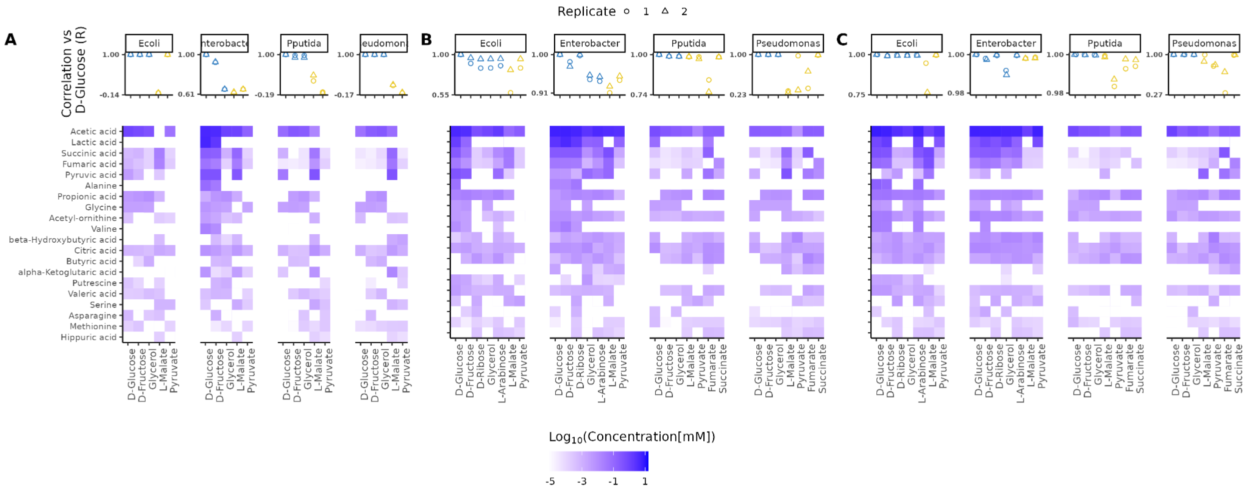


**Supplementary Figure 11: Similarity in secretion profile on different carbon sources.** Heatmaps show the metabolite profile of each strain after 16 hours (A), 28 hours (B), and 48 hours (C) of growth on all carbon sources. Top panel shows Pearson correlation in metabolite abundance compared to D-Glucose. For all strains and timepoints we find that the metabolite profile on other glycolytic carbon sources (blue) is highly correlated to that observed on D-Glucose. Gluconeogenic carbon sources (yellow) display a more idiosyncratic correlation to D-Glucose which varies by strain, by carbon source and by time point.

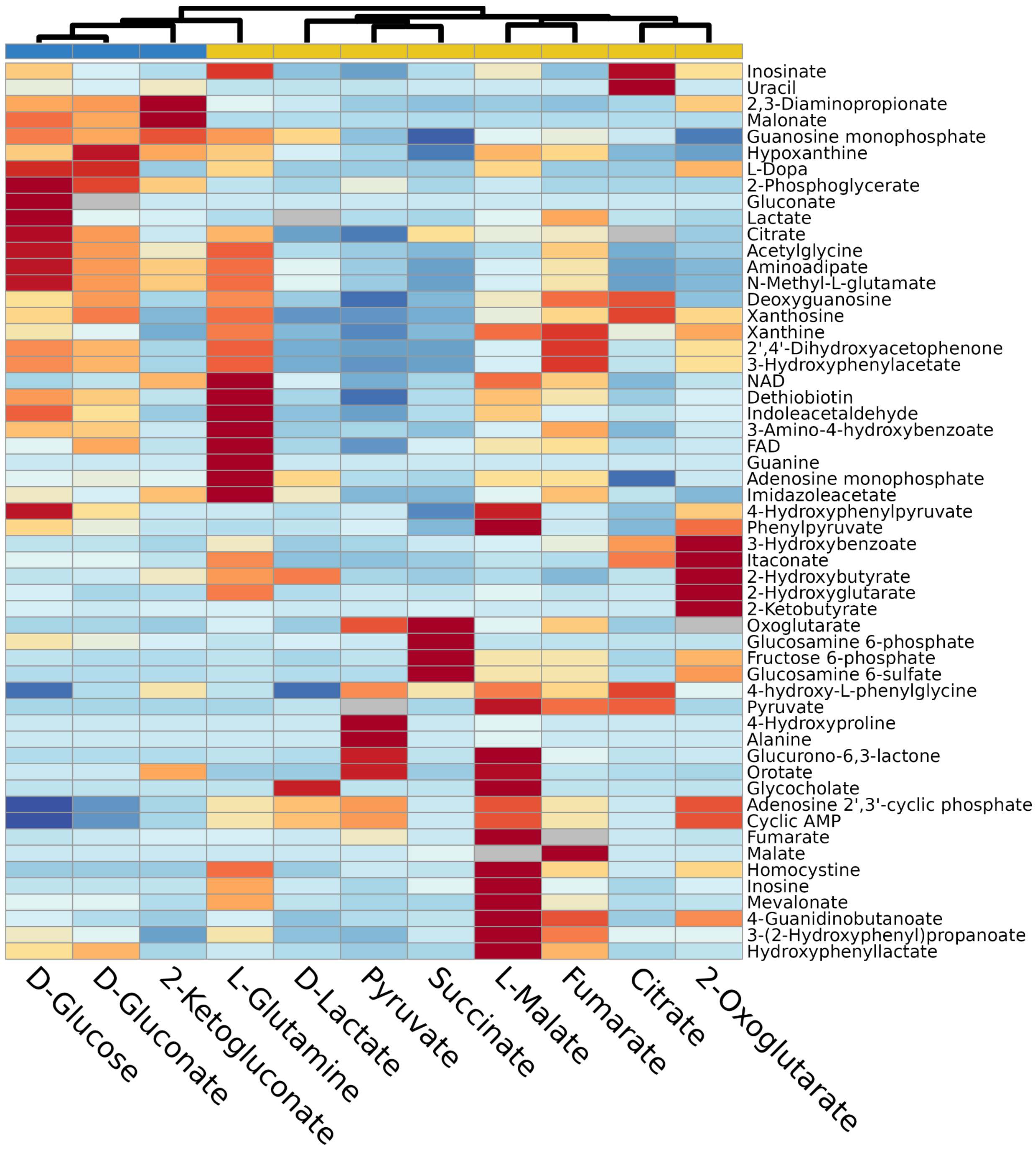


**Supplementary Figure 12: Heatmap shows normalized peak intensity for each metabolite measured using untargeted LC-MS.** In Figure 3C we show the peak intensity averaged across the 3 replicates on each carbon source for the *Klebsiella* strain. Here we show the results for the *Pseudomonas* strain when grown on 11 different carbon sources.

For each metabolite, red corresponds to carbon sources where the metabolite reaches higher abundance, whereas blue corresponds to carbon sources where the metabolite reaches a lower abundance. Carbon sources are hierarchically clustered based on the correlation in by-product profile.

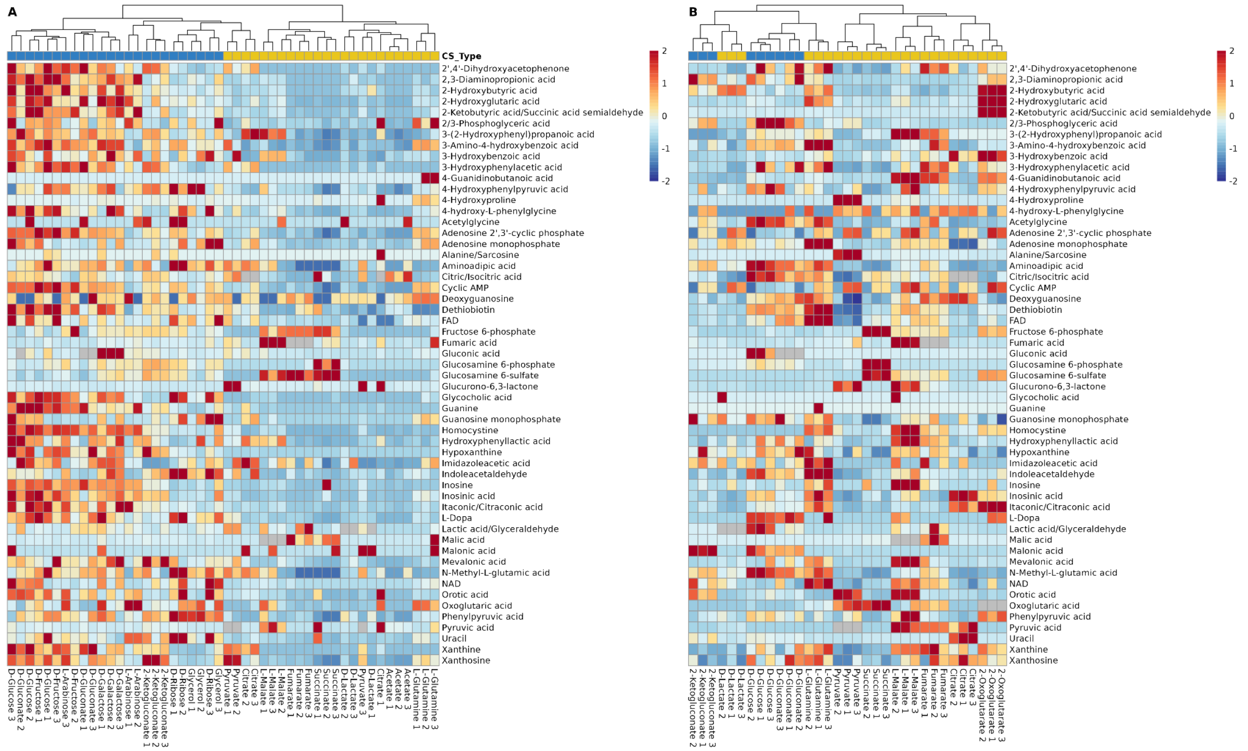


**Supplementary Figure 13: Heatmap shows normalized peak intensity for each metabolite measured using untargeted LC-MS broken down by individual replicate.** In Figures 3C and S11 we cluster carbon sources based on the normalized average peak intensity across the 3 replicates. In panel A we repeat this analysis using all 48 samples for the *Klebsiella* strain (3 replicates of 16 carbon sources). In Panel B we repeat this analysis using all 33 samples for the *Pseudomonas* strain (3 replicates of 11 carbon sources).

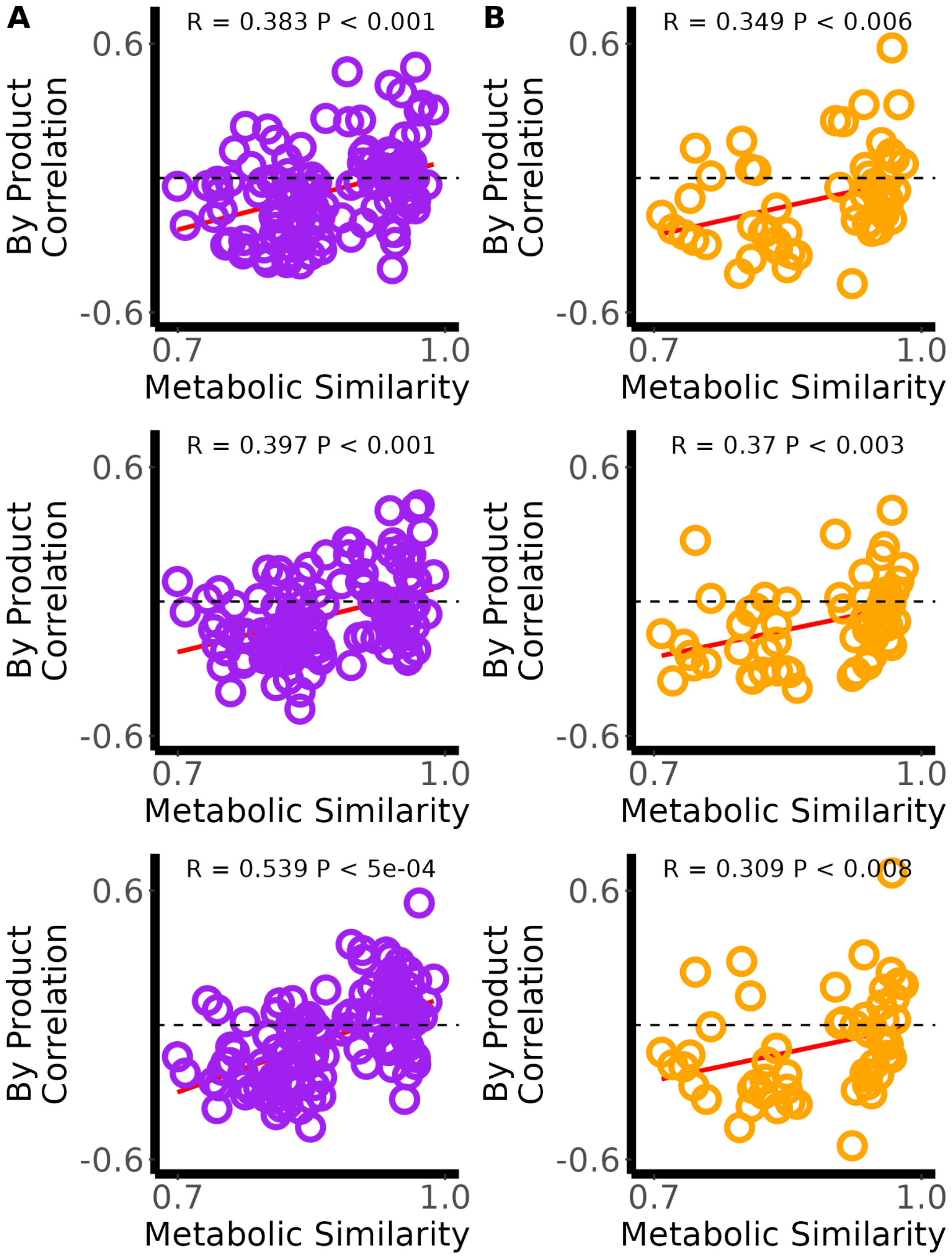
A
R = 0.383 P < 0.001
B
R = 0.349 P < 0.006
R = 0.397 P < 0.001
R = 0.37 P < 0.003
R = 0.539 P < 5e-04
R = 0.309 P < 0.008
By Product Correlation
0.6
-0.6
0.7
1.0
Metabolic Similarity

**Supplementary Figure 14: Relationship between metabolic similarity and by-product profile broken down by individual replicates.** Panel A shows this relationship for the *Klebsiella* strain and Panels B shows this relationship for the *Pseudomonas* strain. In the main figure (Figure 3D-DF) we averaged the peak intensity for each metabolite across the three replicates and calculated the correlation in the normalized average peak intensity. We always observed a positive correlation between metabolic similarity and by-product correlations (Mantel test p<0.01)

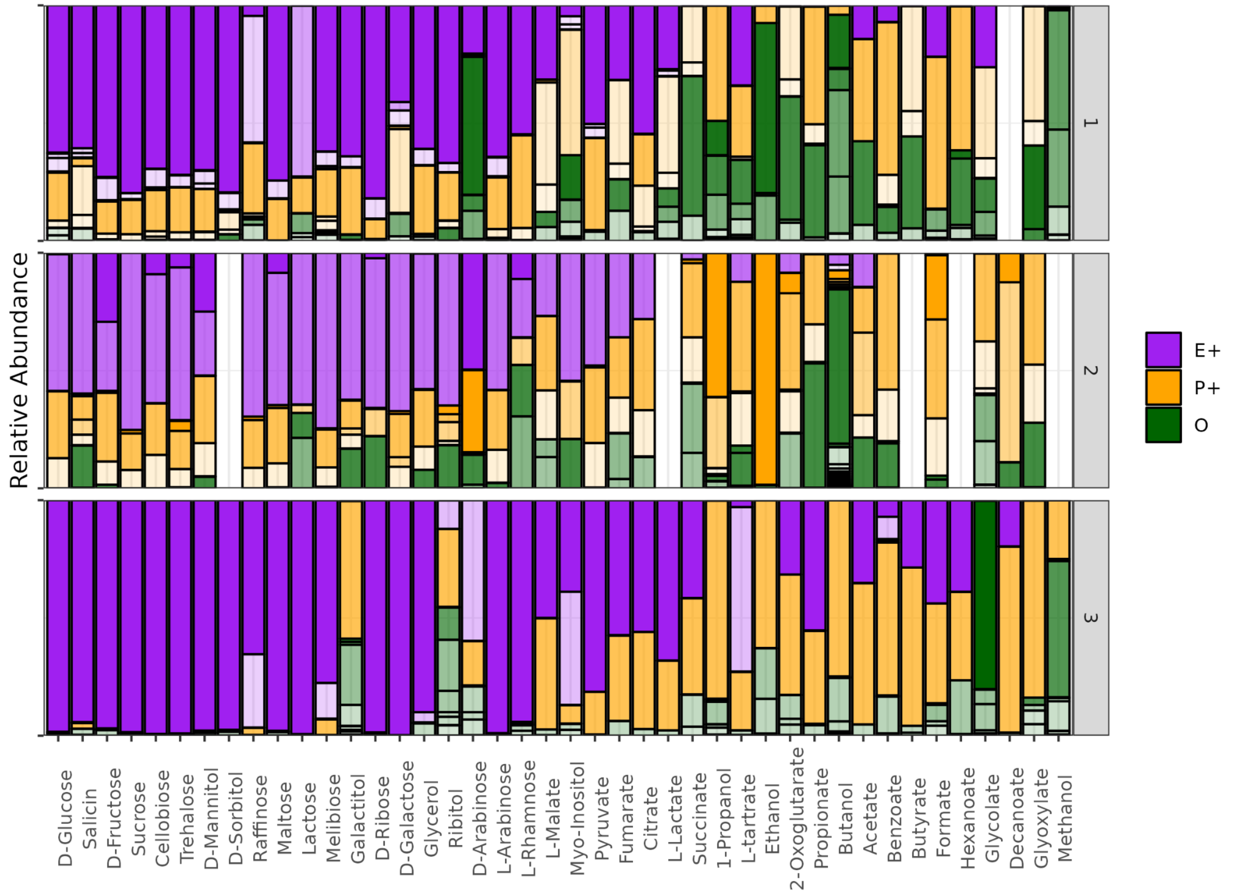


**Supplementary Figure 15 :Composition of communities assembled on one of 42 different carbon sources.** Each row corresponds to a different inoculum and each column corresponds to a different carbon source. Communities are ordered by carbon source metabolic similarity to D-Glucose. Within each community different ESV are shown in different shades of Purple (E+ Strains) , Orange (P+ strains) and Green (Other strains). Missing communities correspond to communities that were not sequenced or had an extremely low sequencing depth (<2000 reads).

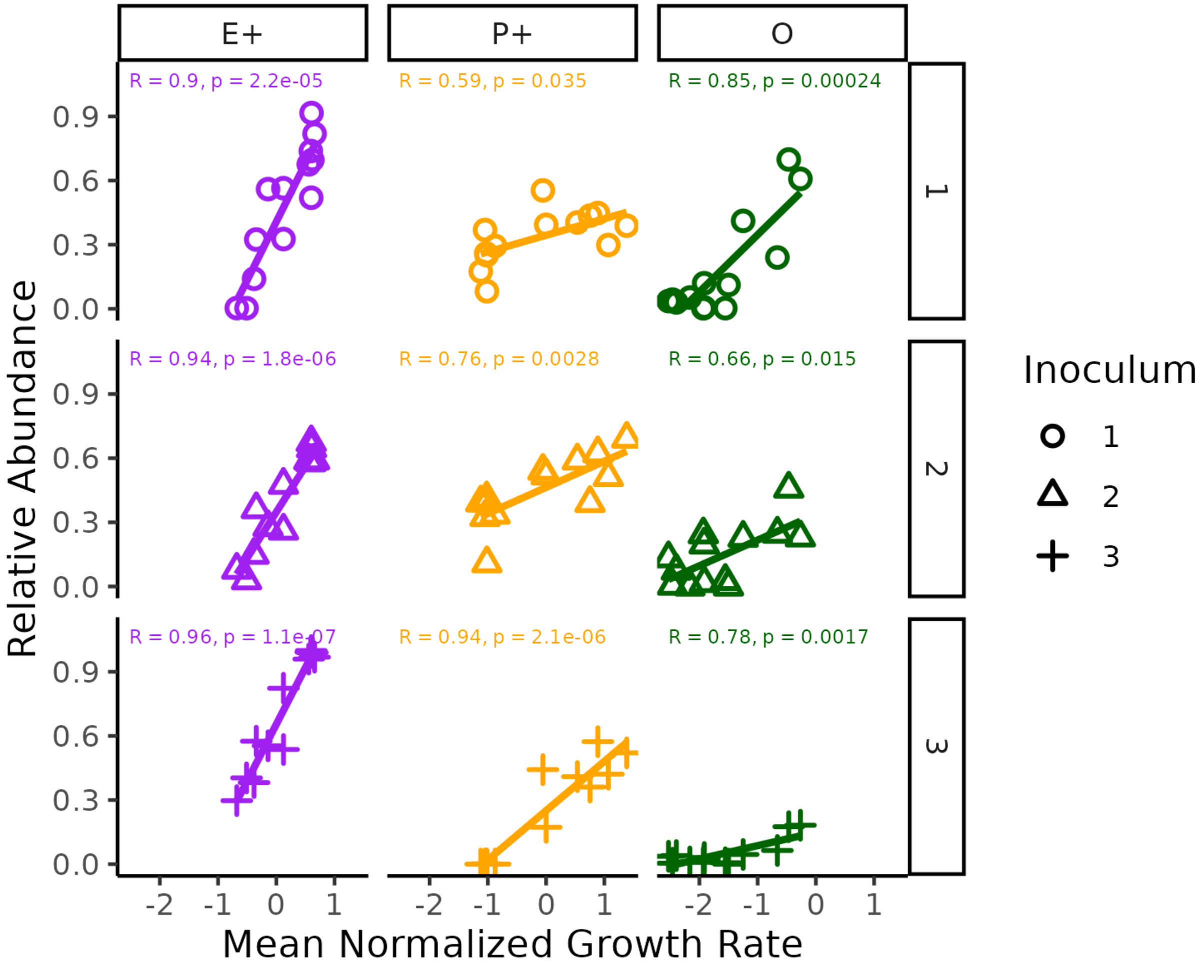


**Supplementary Figure 16. E+ , P+ and O strains reach higher abundances on carbon sources on which members of these clades typically grow faster**. Each data point corresponds to a community assembled on one of 13 carbon sources for which both growth data (Figure 2) and community composition data (Figure 4) was collected. On the X Axis we show the average normalized growth rate of each clade on each carbon source. On the Y axis we show the relative abundance of that clade on the corresponding carbon sources. Different row/shapes correspond to communities assembled from different inoculum.

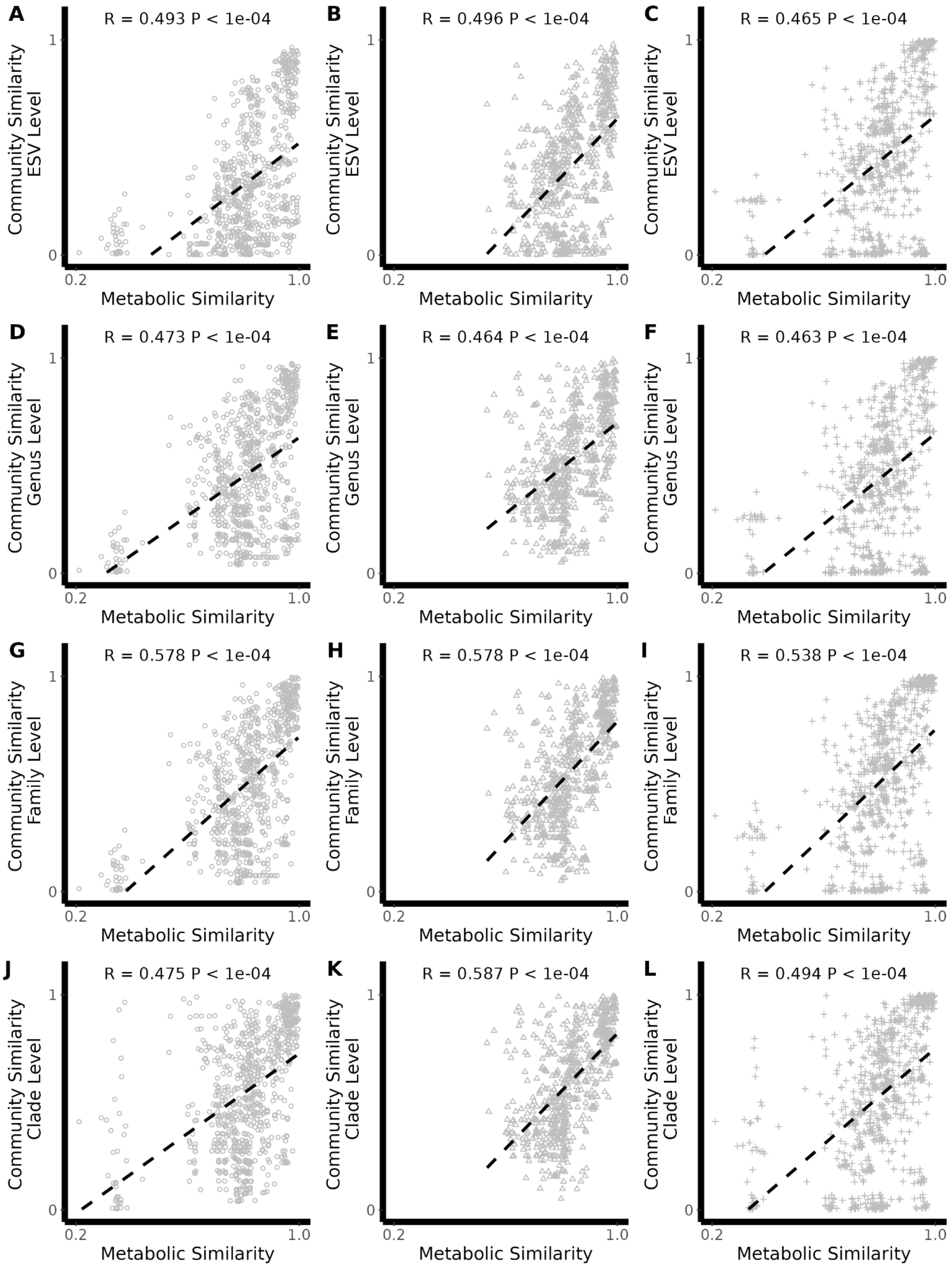
A
R = 0.493 P < 1e-04
B
R = 0.496 P < 1e-04
C
R = 0.465 P < 1e-04
Community Similarity ESV Level
D
R = 0.473 P < 1e-04
E
R = 0.464 P < 1e-04
F
R = 0.463 P < 1e-04
Community Similarity Genus Level
G
R = 0.578 P < 1e-04
H
R = 0.578 P < 1e-04
I
R = 0.538 P < 1e-04
Community Similarity Family Level
J
R = 0.475 P < 1e-04
K
R = 0.587 P < 1e-04
L
R = 0.494 P < 1e-04
Community Similarity Clade Level
Metabolic Similarity
0
1
0.2
1.0

**Supplementary Figure 17: Relationship between substrate metabolic similarity and community similarity (Renkonen similarity) for pairs of communities assembled from the same inoculum at different taxonomic scales**. In Figure 4C- F we show the relationship between metabolic similarity and community similarity at different taxonomic scales for Inoculum 1 (panels A,D,G,J). Similar results are observed on Inoculum 2 (panel B,E,H,K) and Inoculum 3 (panel C,F,I,L). We always observed a positive correlation between metabolic similarity and community similarity (Mantel test p<0.001)

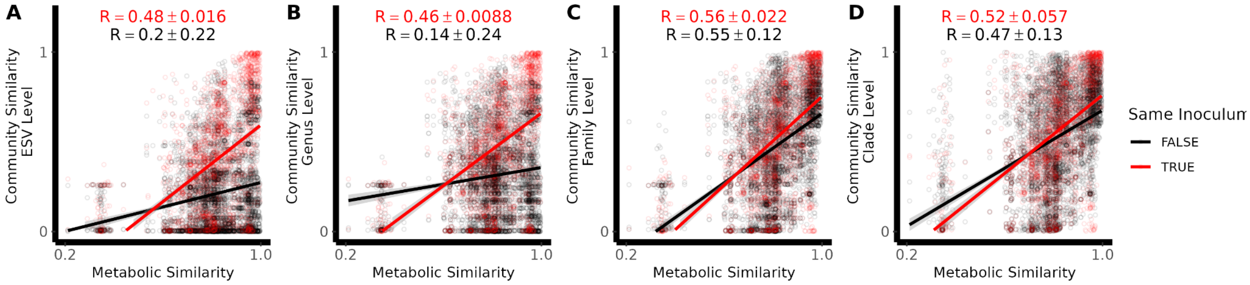


**Supplementary Figure 18:** Pairs of communities assembled from different inoculum have substantially weaker correlations between metabolic similarity and community similarity at the ESV and Genus Level compared to communities assembled from the same inoculum. In contrast, using a different inoculum has a smaller impact at the Family or Clade Level.

:

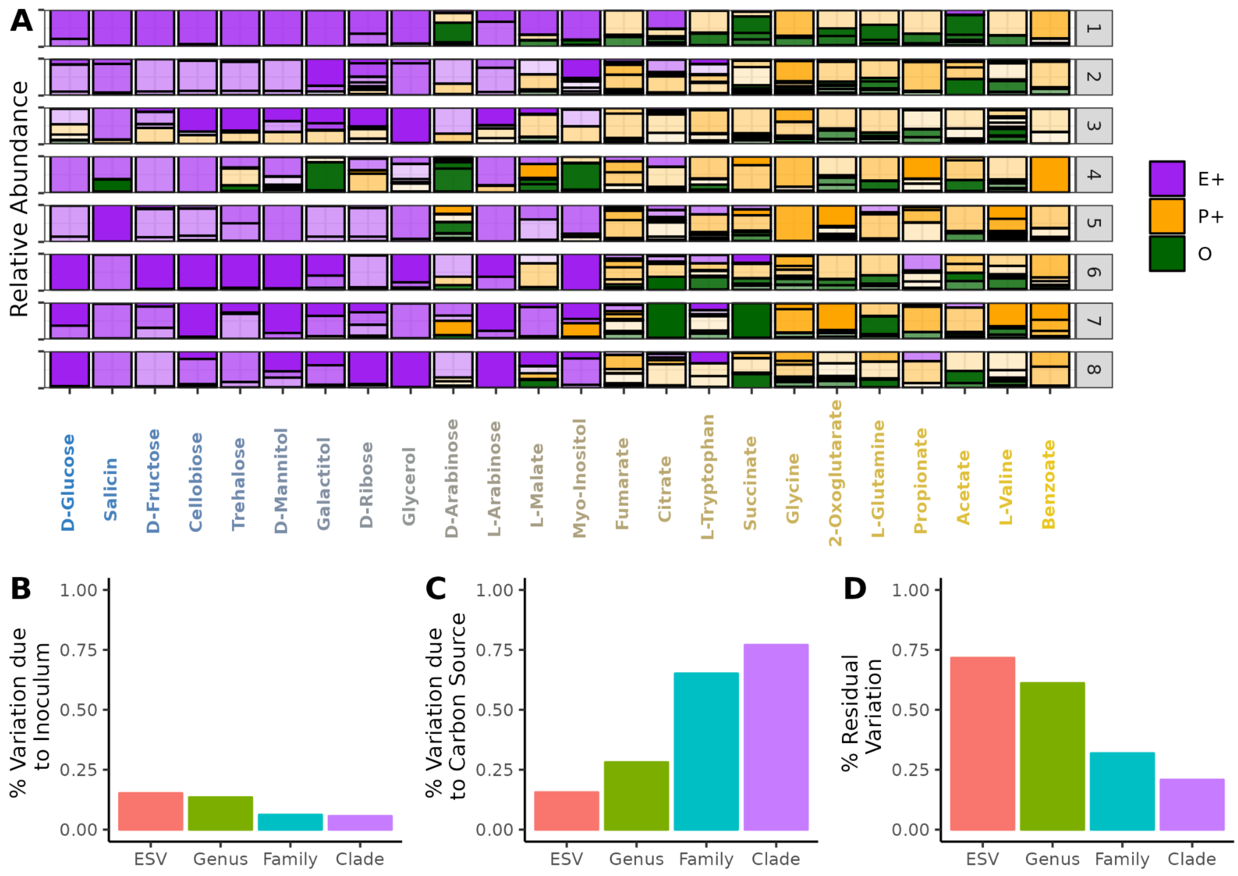


**Supplementary Figure 19: Decomposition of the variation in community composition** (A) Communities from 8 different inocula were assembled on 1 of 24 carbon sources (Estrela al 2022). Using this data we can partition the variation in community composition (beta-diversity) into the variation due to the Inocula, the variation due to Carbon source and the residual variation. In panels B to D we plot the fraction of variation calculated using the *varpart* function in the *vegan* package at different taxonomic scales (Methods,(Legendre 2008)).

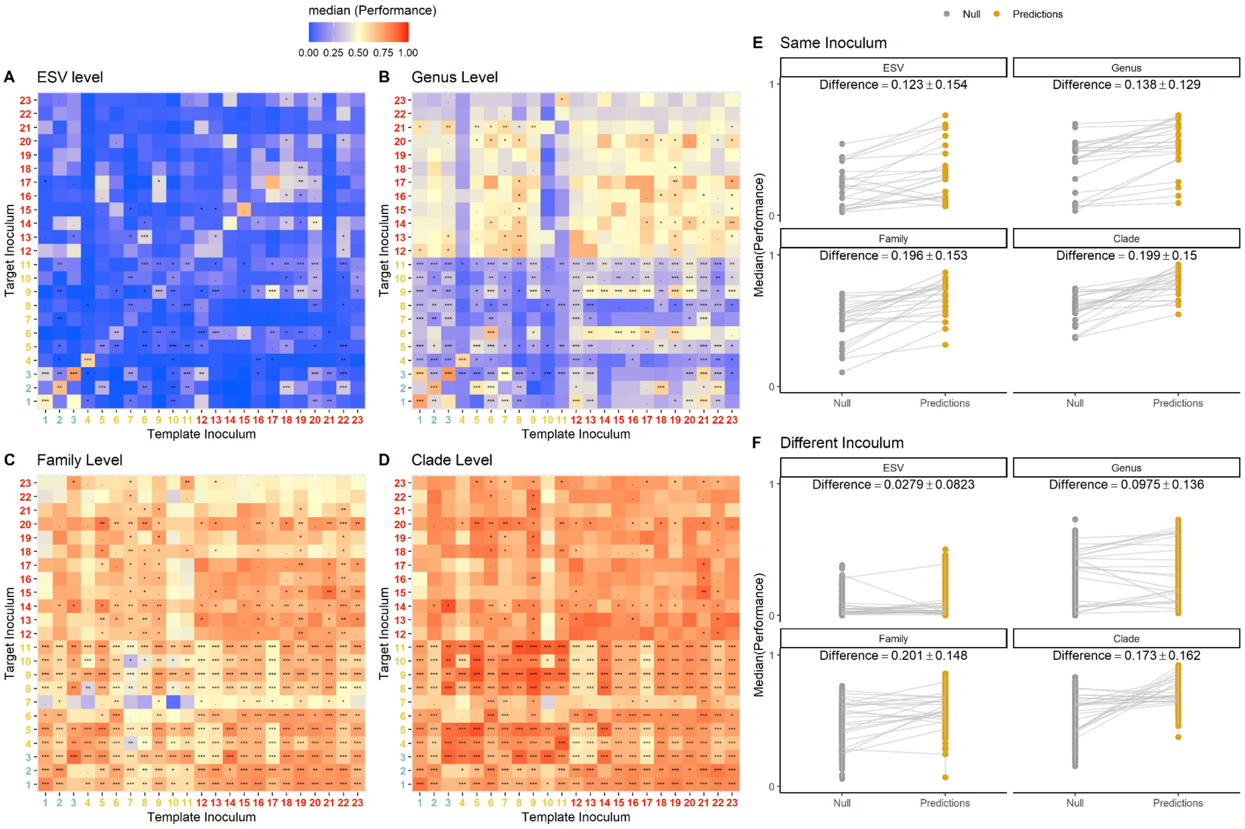


**Supplementary Figure 20: Predictions made using communities assembled from different inoculum generally do better than the null at a coarse-grain taxonomic scale but not at a fine-grain taxonomic scale.** We repeated the “leave one-carbon source out analysis' (shown in Fig 5) using all possible combinations of inoculum from all three experiments as the template or target. Heatmaps show the median performance for each inoculum pair across all carbon sources in the target inoculum. Inocula are colored by whether they come from the 42 carbon source experiment (Fig. 4, Blue) the 24 Carbon source experiment (Fig. S19, Yellow) or the 7 carbon source experiment (Fig. 1,Red). For each pair of inoculum we compared the distribution of performances to the corresponding null distribution using the one-tailed Mann-Whitney U test. Symbol on each grid show the p.value for this test (*** : 0 - 0.001, **: 0.001 - 0.01,*:0.01-0.05 *, .=0.05-0.1). In panel E-F we summarize the results of this analysis. Each data point plots the median performance for a single pair of inocula compared to the median null performance. We show this for predictions made using the same (E) or different inoculum (F). The statistic reports the mean±sd difference between the median performance for predictions and the null

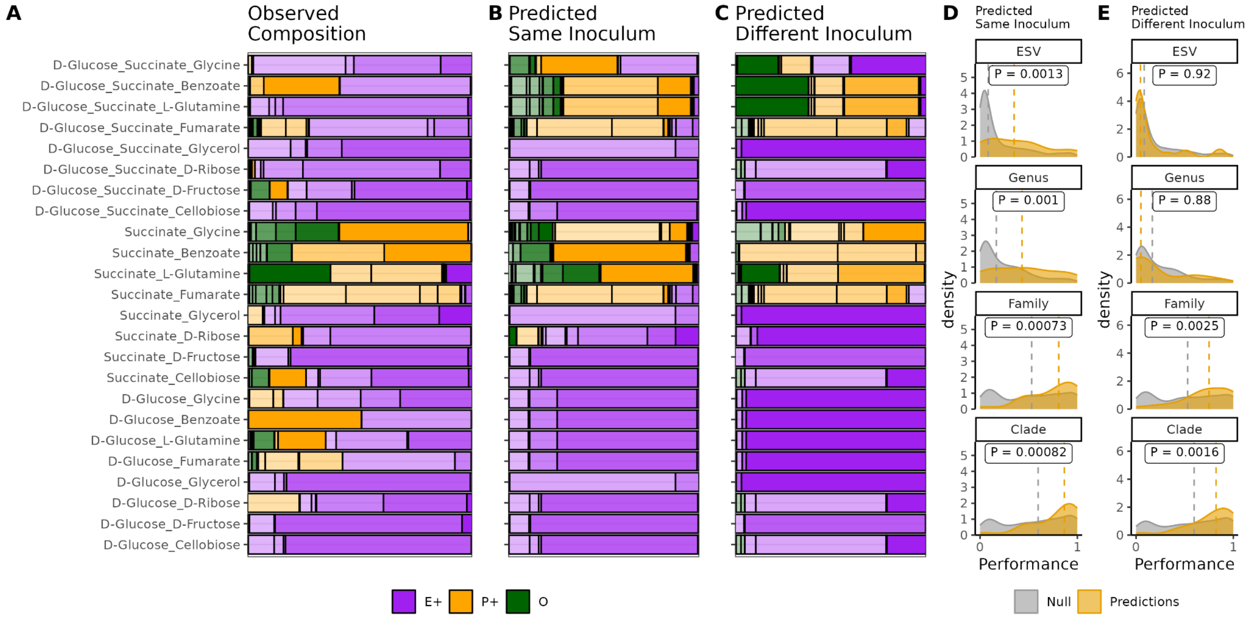


**Supplementary Figure 21: The comparative approach can be extended to predict community in mixed carbon source environments.** (A) Community composition on 18 mixtures of 2 or 3 different carbon sources. (B) Composition predicted in the mixture using communities assembled from the same inoculum on single carbon environments (see Methods). (C) Composition predicted using communities assembled from different inoculum on the single carbon environment. (D) Distribution of performance for predictions made using the same inoculum (yellow) compared to the corresponding null (Grey). (E) Distribution of performance for predictions made using different inoculum (yellow) compared to the corresponding null (Grey).

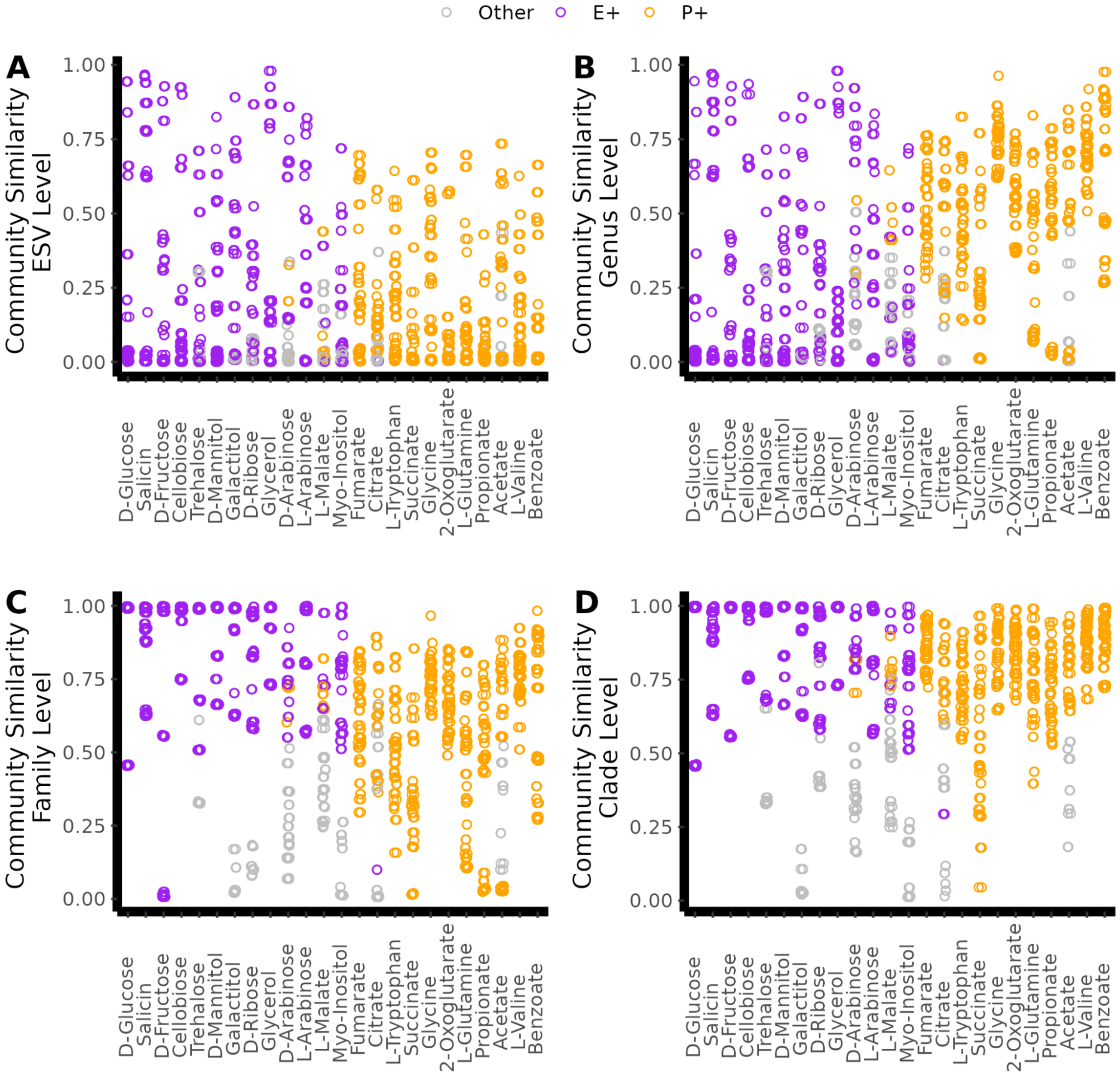


**Supplementary Figure 22.** Communities on which P+ is more abundant are less similar at the family and clade level than communities in which E+ is more abundant. In each panel we plot the Renkonen similarity for pairs of communities assembled from different inocula on each of the different carbon sources. Pairs of communities are coloured based on whether: E+ is more abundant than P+ in both communities (Purple), P+ is more abundant than E+ in both communities (orange) or E+ is more abundant than P+ in one community but not in the other (Grey). Communities in this figure correspond to communities from Figure S19.

**Supplementary Table 1: List of carbon sources considered throughout this study and Additional reactions added to iJO1366 to ensure growth on every carbon.**

| Carbon Source | Biggs ID | Reactions Added |
|---|---|---|
| D-Glucose | glc__D_e | N/A |
| D-Gluconate | glcn_e | N/A |
| Glycerol | glyc_e | N/A |
| D-Fructose | fru_e | N/A |
| L-Arabinose | arab__L_e | N/A |
| D-Galactose | gal_e | N/A |
| D-Ribose | rib__D_e | N/A |
| 2-Ketogluconate | 25dkglcn_e | 25DKGLCNt2rpp 25DKGLCNtex EX_25dkglcn_e |
| Pyruvate | pyr_e | N/A |
| L-Malate | mal__L_e | N/A |
| Citrate | cit_e | N/A |
| Acetate | ac_e | N/A |
| D-Lactate | lac__D_e | N/A |
| Succinate | succ_e | N/A |
| Fumarate | fum_e | N/A |
| L-Glutamine | gln__L_e | N/A |
| L-Phenylalanine | phe__L_e | PPYRDC |
| L-Leucine | leu__L_e | LEUTA OIVD1r MBCOAi MCCC MGCH HMGL |
| 2-Oxoglutarate | akg_e | N/A |
| Sucrose | sucr_e | N/A |
| Melibiose | melib_e | N/A |
| Lactose | lcts_e | N/A |
| Trehalose | tre_e | N/A |
| Cellobiose | cellb_e | EX_cellb_e CLBtex CELBpts BGLA1 |
| Maltose | malt_e | N/A |
| Raffinose | raffin_e | EX_raffin_e RAFFINt2 RAFGH SUCR |

| Carbon Source | Biggs ID | Reactions Added |
|---|---|---|
| D-Sorbitol | sbt__D_e | N/A |
| D-Mannitol | mnl_e | N/A |
| D-Arabinose | arab__D_e | EX_arab__D_e ARAB_Dt ARABDI RBK_Dr |
| L-Rhamnose | rmn_e | N/A |
| Ribitol | rbt_e | N/A |
| Galactitol | galt_e | N/A |
| Myo-Inositol | inost_e | INS2D INSCR |
| Salicin | salcn_e | EX_salcn_e SALCpts S6PG 2HXMPt6 EX_2hxmp_e |
| L-Lactate | lac__L_e | N/A |
| L-tartrate | tartr__L_e | N/A |
| Propionate | ppa_e | EX_ppoh_e PPOHtex PPOHt2pp ALCD3ir |
| Benzoate | bz_e | EX_bz_e BZt BZ12DOX BZDIOLDH CATDOX MUCCYCI MUCLI OXOAEL 3OADPCOAT |
| Butanol | but_e | EX_btoh_e BTOHtex BTOHt2rpp ALCD4y |
| Hexanoate | hxa_e | N/A |
| Decanoate | dca_e | N/A |
| Formate | for_e | N/A |
| Glycolate | glyclt_e | N/A |
| Glyoxylate | glx_e | EX_glx_e GLXt2 |
| Methanol | meoh_e | PRDX |
| Ethanol | etoh_e | N/A |
| 1-Propanol | ppoh_e | N/A |
| Butanol | btoh_e | N/A |
| L-Valine | val__L_e | N/A |
| L-Tryptophan | trp__L_e | N/A |
| Glycine | gly_e | N/A |
